# Why are so many fusogens rod-shaped?

**DOI:** 10.1101/2025.06.30.662463

**Authors:** Ioana C. Butu, Jin Zeng, Dong An, Zachary McDargh, Kunyu Duan, Ben O’Shaughnessy

## Abstract

Molecular fusogens catalyze membrane fusion for many basic biological processes. In eukaryotic cells, SNARE proteins drive membrane fusion for trafficking and for exocytic release in many contexts, from neurotransmission to enzymatic digestion, while other fusogens mediate cell-cell fusion for organ formation, placental development and gamete fusion. Enveloped viruses use glycoprotein fusogens for host cell entry and delivery of the viral genome. Despite this breadth of roles, a structural feature shared by many of these fusogens is their rod shape, conserved across the SNARE superfamily, the class I and II fusogen superfamilies and the class III fusogen family. Here we used highly coarse-grained molecular dynamics (MD) simulations to examine the collective behavior of rod-like fusogens on the microscopically long timescales of physiological membrane fusion. Rod-generated entropic forces maintained a cleared fusion site, squeezed and hemifused the membranes, and then expanded and ruptured the hemifused connection to yield fusion. More fusogens generated higher entropic forces and faster fusion, consistent with electrophysiological measurements at neuronal synapses. The required fusogenic feature was the rod shape, since simulated SNARE complexes, class II EFF-1 fusogens, and model rod-shaped complexes entropically drove fusion along similar pathways, whereas globular complexes failed. Thus, rod-like fusogens are optimally shaped generators of entropic forces that drive membrane fusion. These results suggest a universal rod-based fusion mechanism may have been the evolutionary driver of structural convergence among major classes of eukaryotic and viral fusogens.

**Significance:** Large classes of eukaryotic and viral membrane fusion complexes are rod-shaped, a common structural feature despite a multiplicity of functions, from neurotransmission to organogenesis to cell entry. Here we used coarse-grained molecular dynamics to access the millisecond timescales of physiological fusion. Simulations revealed a universal mechanism whereby, regardless of structural details, rod-shaped fusogens generate entropic forces that drive fusion. This suggests the mechanism was a major evolutionary driver of structural convergence and multiple transmissions between eukaryotic hosts and viruses. Rod-mediated fusion is faster with more fusogens, explaining why release at presynaptic nerve terminals is enhanced or suppressed depending on the number of active SNARE fusogens. Thus, modulation of SNARE numbers and fusogenicities may represent a significant mechanism of synaptic plasticity.

## Introduction

Compartmentalization by lipid membranes is essential to all forms of life (1), but for many fundamental processes these compartments need to be transiently opened and merged with other membrane-enclosed compartments. For this purpose cells use specialized membrane fusion machineries whose core consists of fusogens, molecular fusion proteins that perform the membrane fusion step. In eukaryotes, proteins of the SNARE superfamily mediate membrane fusion reactions for intracellular trafficking (2) and for exocytosis in which intracellular vesicles fuse with the plasma membrane to mediate basic processes, from neurotransmitter release at neuronal synapses (3), to hormone release (4), to digestive enzyme secretion (5).

Other fusogens drive cell-cell fusion for developmental processes, including organ formation, placental development and gamete fusion. Syncytins are cell-cell fusogens associated with placental morphogenesis, identified in many mammals (6). In *C. elegans* and other nematodes, the EFF-1 fusogens are involved in epithelial organ sculpting (7), phagosome sealing (8) and neuronal dendritic tree maintenance (9), while AFF-1 fusogens mediate vulval development (10). Fusion of myoblast muscle precursor cells in vertebrates requires the muscle-specific fusogens Myomaker and Myomerger (11). HAP2 is an ancient gamete fusogen found in many eukaryotes (12). Membrane fusion is also central to the life cycle of pathogens such as enveloped viruses, whose glycoprotein fusogens merge the viral and host membranes for host cell entry and delivery of the viral genome (13).

Most of these fusogens are rod-like, a structural similarity across a remarkably broad range of functions and contexts (Fig. 1). (Here, we refer to the folded complex that immediately precedes and drives fusion.) SNARE complexes, assembled from cognate SNAREs associated with each of the fusing membranes, are ∼ 12 nm long, ∼ 2 nm wide coiled-coil rod-like bundles of four 𝛼-helices, attached by ∼ 10-residue linker domains (LDs) to transmembrane domains (TMDs) that anchor the complex to the vesicle and target membranes (14, 15). This structure is similar to that of class I viral fusogens, as noted long ago (16). Class I fusogens are also 𝛼-helical rods, including those of influenza, SARS CoV-2, HIV and Ebola viruses (17). Following a transition from the pre-fusion to the extended intermediate state (18), the viral glycoprotein folds into a rod-like “hairpin” *trans* complex consisting of three helices in a coiled-coil backbone together with three shorter antiparallel helices (17). A similar six-helix rod-like structure is adopted by the eukaryotic syncytins, derived from the *env* genes of ancient retroviruses that integrated into host germline cells (6). Accordingly, the class I superfamily has been defined (19), encompassing both viral fusogens and eukaryotic syncytins. All superfamily members have the characteristic six-helix rod-like shape.

**Figure 1.**
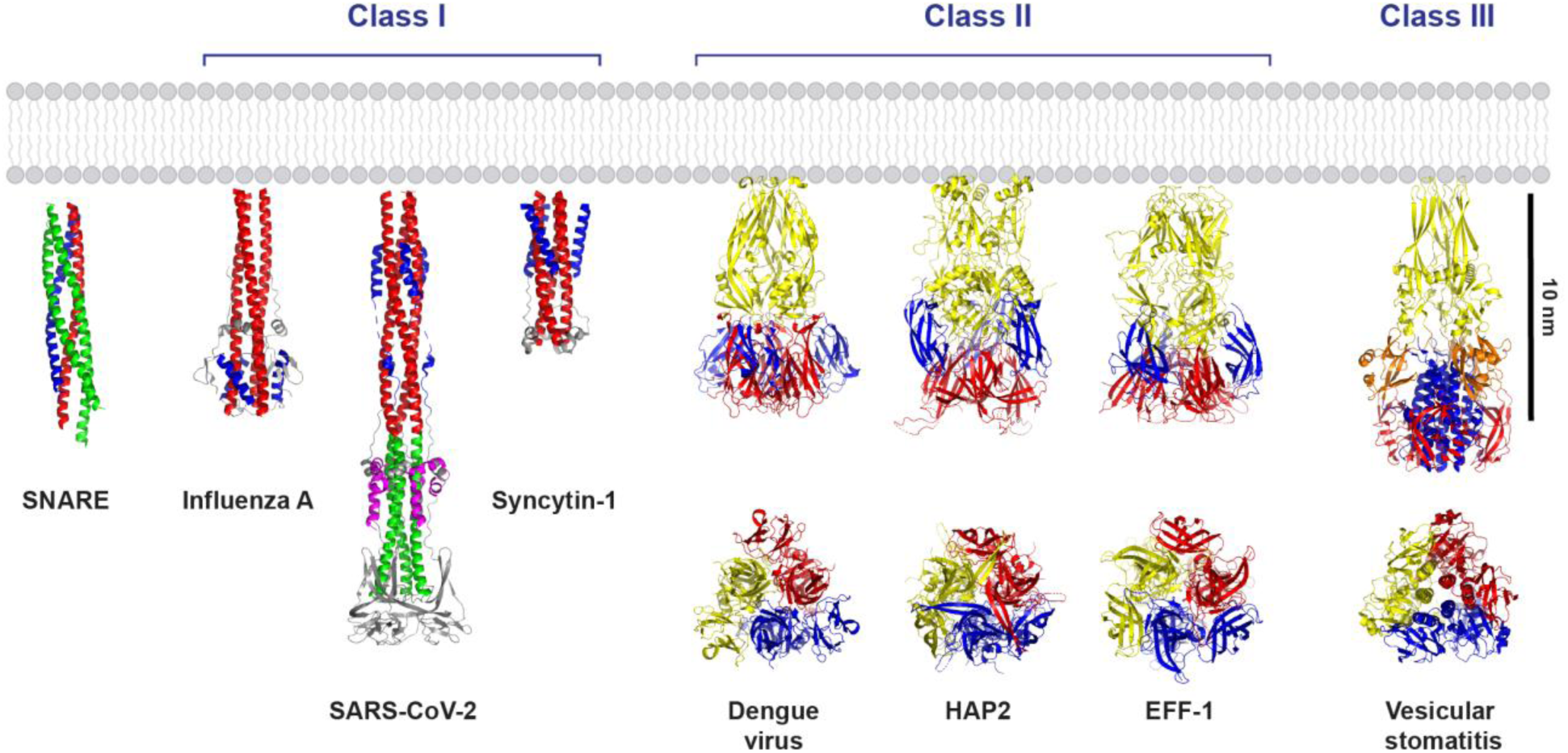
Post-fusion crystal structures of eukaryotic and viral fusogens. The neuronal SNARE complex (PDB: 3HD7) is a coiled coil rod of four 𝛼-helices. Class I superfamily fusogens are trimers, with an extended three-helix coiled coil backbone (red/green) and three shorter anti-parallel helices (blue/magenta) completing a six-helix bundle in one or more portions of the rod. Members include eukaryotic fusogens such as the human syncytin-1 (PDB: 6RX1) cell-cell fusogen involved in placental morphogenesis. Class II superfamily fusogens have high β-sheet content and are also trimeric in the post-fusion state, each protomer having one elongated (yellow) and two globular (red, blue) domains. Examples include the *C. reinhardtii* HAP2 gamete fusogen (PDB: 6DBS) and the *C-elegans* EFF-1 (PDB: 4OJC) cell-cell fusogen involved in organ development. Class III viral fusogens combine structural features of classes I and II. Domains I-IV are colored red, blue, orange, and yellow, respectively. End-on views (classes II and III) depict the three protomers red, blue, and yellow, respectively.

The rod-like structural theme continues into class II and class III fusogens, although the rods are considerably more bulky (Fig.1). The class II superfamily includes fusogens of viruses such as dengue fever, yellow fever and rubella, and eukaryotic members such as the EFF-1 cell-cell fusogens and the HAP2 gamete fusogens, both thought to be endogenous retrovirus fusogens descended from ancient class II retroviruses (20). Class II fusogens are rod-like trimers with high 𝛽-sheet content. Typically longer than ∼ 10 nm, an elongated core of finger-like domains leads to a broader membrane-distal head where additional globular domains pack against the core. Finally, class III viral fusogens combine 𝛼-helical and 𝛽-sheet features of classes I and II, with a trimeric rod-like structure featuring an elongated 𝛽-sheet base, a central coiled-coil, and additional 𝛽-sheet-rich domains (17). Members include the fusogens of vesicular stomatitis virus and herpes simplex virus 1.

In summary, rod-like structure is a common feature of eukaryotic SNAREs, numerous eukaryotic cell-cell fusogens, and class I, II and III viral fusogens. Other fusogens with different structures do indeed exist, such as the dynamin-like mitochondrial fusogens mitofusin and optic atrophy 1 (21), the muscle fusogens Myomaker and Myomerger (11), and the class IV viral fusion-associated small transmembrane (FAST) cell-cell fusogens encoded by nonenveloped reoviruses (22). Nonetheless, the abundance of rod-like eukaryotic and viral fusogens raises the possibility that the rod-like structure is mechanistically significant.

Indeed, the mechanisms of membrane fusion are far from established. The most thoroughly studied example is SNARE-driven fusion. A common view is that fusion is driven by the energy released as the vesicle v-SNARE zippers onto the target membrane t-SNAREs, forming the SNARE complex (23). However, the hypothetical mechanism that would transduce complexation energy into membrane energy is unknown. One proposal is that zippering energy is stored as linker domain (LD) bending energy (24, 25), but the uncomplexed LDs appear unstructured (26, 27) and incapable of such energy storage. Another difficulty with zippering energy hypotheses is that zippering likely occurs on 𝜇s timescales (28), whereas electrophysiological studies suggest membrane fusion and neurotransmitter release at neuronal synapses require ms timescales (29–31). A puzzling aspect is that the number of SNAREs required for fusion as suggested by different experimental studies is highly variable, two to eight *in vivo* (32, 33) and an even broader range *in vitro* (34, 35).

Molecular dynamics (MD) simulations have provided powerful insights into the pathway and mechanism of membrane fusion driven by SNAREs and other fusogens. However, the ms or greater timescales of physiological fusion (29–31) are far beyond the reach of atomistic MD or MD simulations using the coarse-grained Martini force-field. In Martini simulations, SNAREs fused 20 nm pure PE lipid vesicles (36), or nanodiscs were fused with a planar membrane (37). These groundbreaking studies revealed inverted micelle (38) and hemifusion intermediates in which only the outer proximal bilayer leaflets are fused, consistent with many *in vitro* (39, 40) and *in vivo* (41–44) experimental studies. Atomistic simulations drove SNARE-mediated fusion between a 24 nm vesicle and planar membrane (45). However, these Martini or atomistic studies drove fusion on computationally accessible µs timescales, far less than physiological times, likely due to the small vesicle sizes and lipid compositions used (36), the assumption of structured LDs (36, 37), or the imposed constraint that the LDs be fully zippered (45). A hybrid all-atom-Martini approach simulated influenza hemagglutinin-mediated fusion of small 15 and 30 nm liposomes with planar bilayers on μs timescales (46). To study SNARE-mediated fusion we previously used an alternative highly coarse-grained approach that captured ms fusion timescales (47–49). In contrast to a hypothesized zippering energy mechanism, fusion in these simulations was driven by entropic forces among the rod-like SNARE complexes and membranes.

Why do eukaryotes and viruses use rods to fuse membranes? Here, we conclude that the essential fusogenic feature is the ability of rods to generate large entropic forces. We find that structural differences among different rod-like fusogens modify details of the fusion pathway, but they use the same essential fusion mechanism. To deal with the timescale problem, we use MD simulations with highly coarse-grained phospholipid and fusogen representations to capture the cooperative effects of multiple fusogens and access the ms timescales of physiological fusion under physiological conditions (vesicle size, temperature, membrane tension). Regardless of structural details, different rod-like fusogens used entropic rod-rod and rod-membrane forces to maintain a cleared fusion site, to squeeze and hemifuse the membranes, and finally to expand and rupture the hemifused connection to yield fusion. More fusogens generated higher entropic forces and faster fusion, consistent with electrophysiological measurements at neuronal synapses (50, 51). The required fusogenic feature was the rod shape, as model rod-shaped complexes were equally efficient entropic force generators and fusogens as coarse-grained SNARE complexes or EFF-1 fusogens, whereas globular complexes accumulated at the fusion site and failed to drive fusion. Thus, our results suggest that a universal rod-based fusion mechanism may have been a key evolutionary driver of structural convergence within major classes of eukaryotic and viral fusogens.

## Results

### Model

#### Coarse-grained representation of phospholipids

Each lipid has one hydrophilic head bead H and three hydrophobic tail beads T, interacting via the Cooke-Deserno force field (52–54). H-H and H-T interactions are repulsive and hard core, while T-T interactions are repulsive with an attractive tail and well depth 𝜖 = 0.6 𝑘_𝐵_𝑇, representing hydrophobic attraction between lipid tails (52–54). The characteristic hard core diameter was set to 𝜎 = 0.88 nm by equating the simulated bilayer thickness to a typical value of 5 nm for biological membranes (55). We use a simulation time step Δ𝑡 =0.068 ns by equating the simulated lateral lipid diffusivity 8.8 × 10^−5^ 𝜎^2^/Δ𝑡 to the experimental value of 1μm^2^s^−1^in a mixed lipid membrane (DOPC/SM/CHOL) (56).

#### SNARE complex

The coarse-grained model of the neuronal SNARE complex is based on the crystal structure (PDB: 3HD7) (15) as described in (49), adapted from our previous model (47, 48) (Fig. 2A). The 16 layer SNARE complex is a coiled-coil of four 𝛼-helices, two from SNAP-25, and one each from VAMP and syntaxin (Stx) (14, 15), Fig. 1. Groups of four contiguous residues along each helix are mapped to a coarse-grained bead, giving ∼ one layer per bead.

**Figure 2.**
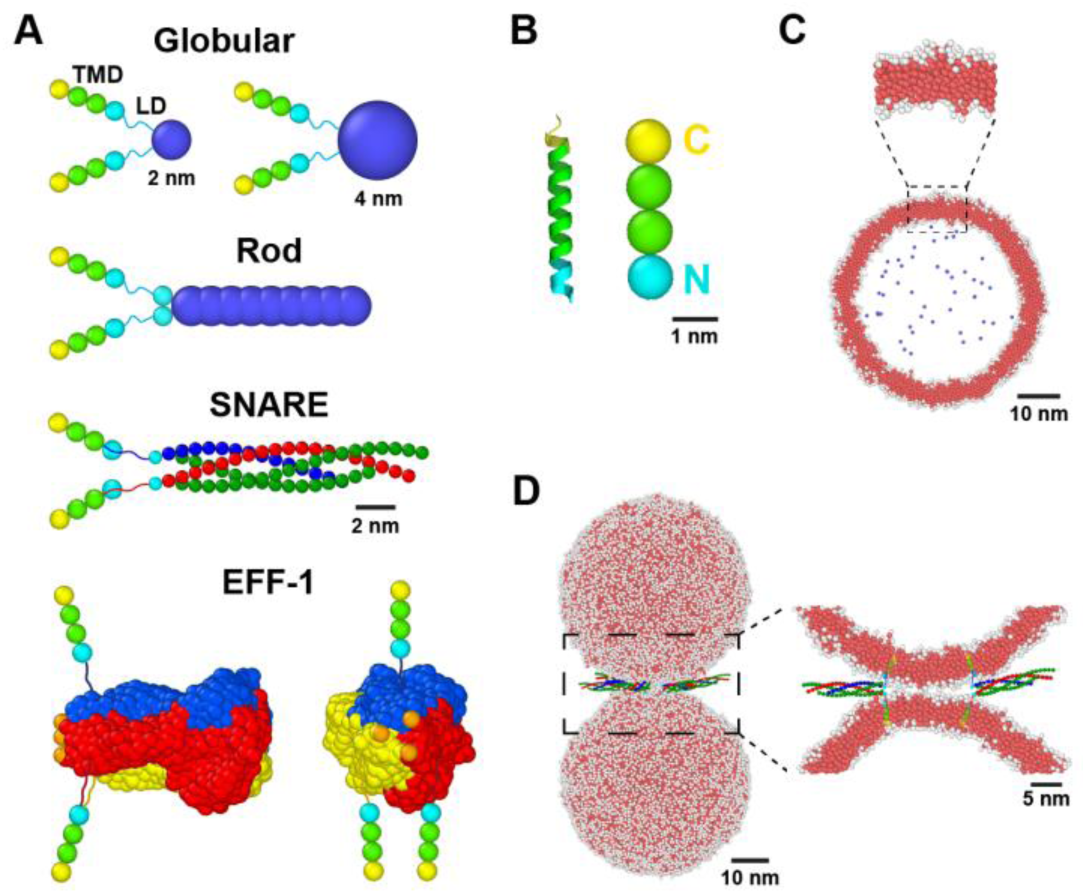
Coarse-grained simulations of membrane fusion. (A) Coarse-grained representations of globular, model rod-like, SNARE, and EFF-1 fusogens. The SNARE complex is built from coarse-grained 𝛼-helices. In the EFF-1 trimer, orange beads represent the acidic patch. (B) Left: syntaxin transmembrane domain (TMD) and the polybasic juxtamembrane end of the linker domain (LD) (cyan) (PDB: 3HD7). Right: coarse-grained TMD for all fusogens: hydrophobic beads (green), a hydrophilic staple bead (yellow), and a hydrophilic LD staple bead (cyan). For EFF-1, the two TMD staple beads are neutral. (C) Cross-section of a simulated vesicle. Coarse-grained phospholipids have one hydrophilic head bead (white) and three hydrophobic tail beads (red). Membrane tension is controlled by fictitious solvent beads (blue) in the vesicle lumen. (D) Initially, two vesicles are bridged by variable numbers of fusogens in trans configuration.

#### Model rod-like fusogen

We constructed a minimal rod-shaped fusogen with similar dimensions to the SNARE complex, a nine-bead rigid body 10 nm long and 2 nm wide (Fig. 2A) with hard core repulsive bead interactions. The **non-rod-like globular fusogen** is identical, but the rod domain is replaced by a single bead (Fig. 2A).

#### EFF-1

The coarse-grained EFF-1 class II cell-cell fusogen (Fig. 2A) represents the folded trimer directly preceding fusion. Its shape is that of the solvent-excluded surface taken from the crystal structure (PDB: 4OJC) (57) (Fig. 1). Three beads (one orange bead per protomer) at the membrane-proximal end of EFF-1 represent negatively charged loops that, together with adjacent residues, form a strong acidic patch thought to interact with polar lipid head groups (57, 58).

#### Transmembrane domains (TMDs)

Simulated TMDs are coarse-grained versions of the 𝛼-helical TMDs of VAMP and Stx, represented as rigid bodies 3 nm long and 1 nm wide, with three repulsive hard core beads (Fig. 2B). A fourth bead represents a staple (see below) and belongs to the linker domain (LD). The central two beads have attractive hydrophobic interactions with the central beads of other TMDs and with lipid tail beads based on reported hydrophobicities (15). TMD-lipid attractions were chosen to prevent TMD pullout. Now the lysine- and arginine-rich polybasic regions at the cytosolic LD-TMD interfaces of VAMP and Stx (59–61) and the negatively charged carboxylic acid groups at the luminal TMD ends (62) are thought to staple the TMD ends to the lipid head groups. Accordingly, at the TMD ends we used staple beads interacting attractively with lipid head beads. EFF-1 TMD end beads are neutral. Since EFF-1 assembles from three identical monomers anchored in opposing membranes and interacting in *trans,* in simulations EFF-1 has one TMD embedded in one vesicle and two TMDs in the other (Fig. 2A).

#### Linker domains (LDs)

For SNAREs, the LDs connecting TMDs to the rod complex are assumed uncomplexed and unstructured, consistent with the LD tensions ∼ 21 pN emerging from our simulations, well above the experimental LD unzippering threshold ∼ 12 pN (23) (Fig. 2A). The LDs are represented as worm-like chains (23) with contour length 3.65 nm (10 residues, each of 0.365 nm uncomplexed length) and persistence length 0.5 nm, typical of unstructured polypeptides (63, 64). These values minimize the LD tension required for seven SNAREs to drive fusion on physiological ms timescales (49). The same LD system is used for EFF-1, where the unstructured LD consists of the ten residues upstream of the TMD in the stem of each of the three monomers. For our model rod-like and non-rod-like globular fusogens, for simplicity we assumed a fixed LD tension of 18 pN down to 0.1 nm LD length, similar to the 21 pN for SNARE LDs. Below 0.1 nm the force decays linearly to zero (23).

#### Simulations and vesicle membrane tension

Fusogens bridge two 50 nm diameter synaptic vesicle-sized vesicles in a 88 nm x 88 nm x 123 nm box with periodic boundary conditions (Fig. 2D). Membrane tension 𝛾_ves_was set by pressure from a fictitious gas of ghost beads within the vesicle (see SI). We used 𝛾_ves_ = 0.05 pN/nm (65) except for brute force simulations where 𝛾_ves_ = 1 pN/nm. Simulations were run using the HOOMD-blue toolkit in the NVT ensemble with a Langevin thermostat (66, 67).

### Brute force drives membrane fusion and post-fusion intralumenal vesicles

We first tried to fuse membranes by simply pressing them together. A previous simulation study fused two ∼ 30 nm diameter coarse-grained lipid vesicles (68) by applying forces to every lipid bead which were released at the onset of lipid exchange. Here, a constant force was applied to every bead and the vesicles were laterally constrained to prevent sliding past one another (Fig. S1). Pressing two 50 nm vesicles together generated a large contact zone of radius ∼ 20 nm (Fig. S2A). For high forces a transient localized hemifusion connection, a stalk, occasionally developed at the contact zone edge with lifetime ∼ 1-10 μs before disassembling. Forces exceeding ∼ 675 pN fused the membranes via a short-lived leaky intermediate: one or more of the stalks elongated along the contact zone edge (Figs. 3A, S2B), a simple pore (a hole in a single bilayer) then developed in each vesicle adjacent to the growing stalk, and the stalk then encircled the two simple pores to create a fusion pore (69). Interestingly, the forces required for fusion correspond to a pressure ∼ 5 atm over the membrane-membrane interface, similar to but somewhat lower than the minimum pressure of ∼ 10 atm to irreversibly hemifuse DMPC bilayers on soft supports (70).

**Figure 3.**
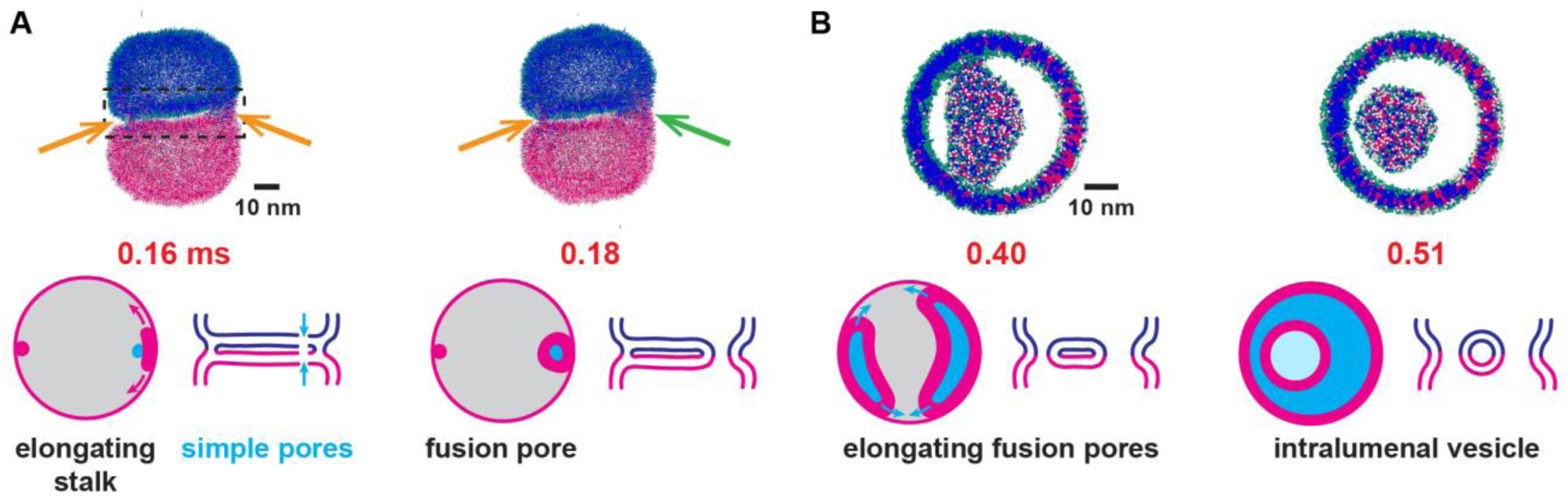
Fusion by brute force. Two 50 nm vesicles are pressed together by a 675 pN body force, laterally stabilized by a potential. Lipid head beads, white or green; tail beads, magenta or blue. (A) By 0.16 ms, two hemifusion stalks (orange arrows) have nucleated at the edge of a large flat membrane interface. The right stalk has elongated along the interface edge and a simple pore formed in each vesicle (blue arrows) (see also Fig. S2). By 0.18 ms the right stalk has encircled and sealed the simple pores, creating a non-leaky fusion pore (green arrow). (B) Orthographic projections of contents of dashed box in (A). The left stalk matured into a fusion pore and both fusion pores elongated, creating an intralumenal membrane tube. By 0.51 ms the fusion pores have merged, releasing an intralumenal vesicle.

Following fusion, the fusion complex underwent topological changes. Typically, several fusion pores created as above subsequently expanded along the contact zone edge and merged to form elongated slits, creating an intralumenal membrane tube (Figs. 3B, S2C, D). Finally, the slits merged, releasing an intralumenal vesicle (Fig. 3B). Occasionally, tubes severed at one end only and the intralumenal tube merged back into the outer membrane instead of forming a vesicle. Sometimes multiple interconnected tubes formed in the intralumenal cavity.

In summary, a brute force approach to fusion generates a large membrane interface whose high curvature boundaries promote lipid tail exposure, activating hemifusion and fusion events at the boundary and a post-fusion intralumenal vesicle. Interestingly, this pathway is similar to the fusion pathway for micron-sized yeast vacuoles (71). We return to this point in Discussion.

### Non-rod-like complexes fail to catalyze fusion

Cells use specialized molecular fusogens. How do the mechanisms compare to fusion driven by simple force? We first simulated globular fusogens of diameter 2 nm, of order the thickness of rod-like SNARE complexes, Fig. 2A. Other simulations used smaller or larger globular domains (1, 3, and 4 nm diameter). In all cases, the LDs pulled the vesicles together and the fusogens aggregated at the vesicle-vesicle contact point for most of the simulation, blocking hemifusion and fusion (Figs. 4A, S3, Movies S1, S2). No hemifusion or fusion occurred in a total of ∼ 30 ms simulation time for 2 nm fusogens or ∼ 15 ms for the other sizes. The smallest 1 nm fusogens occasionally provoked thin membrane connections that rapidly severed within ∼ 1-10 μs.

**Figure 4.**
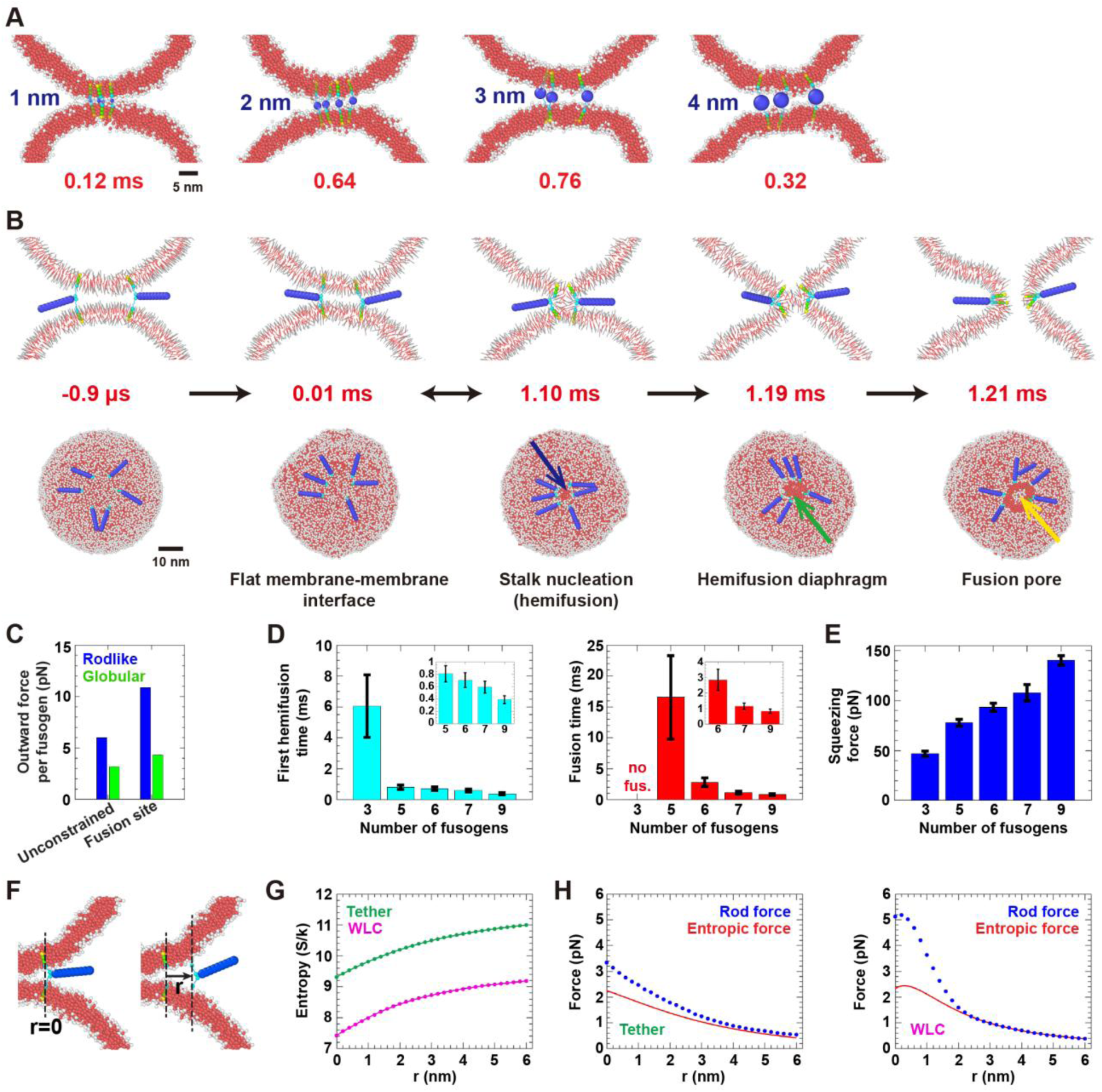
Model rod-like fusogens generate entropic forces that drive fusion but non-rod-like globular complexes fail as fusogens. (A) Globular complexes of four diameters accumulated between the vesicles and failed to hemifuse or fuse membranes (total running times ∼ 14 ms for 1, 3 and 4 nm and ∼ 28 ms for 2 nm). See also Fig. S3. Cross-sections ∼ 5 nm thick. (B) Following 1 μs of equilibration, thermal collisions pushed model rod-like fusogens outwards, maintaining a flat membrane interface with rods at the edge. The fusogens nucleate a stalk, expand the stalk into a hemifusion diaphragm (HD) and rupture the HD for fusion (see also Fig. S4). Often fusion is preceded by reversible stalk episodes (double arrows). Top row: cross-sections, lipids in line segment representation. (C) Time-averaged radial force per fusogen for normal simulations (unconstrained, 𝑛 = 10 simulations) and with TMDs constrained in a 3 nm radius ring at the fusion site (n = 5 simulations). For SDs, see SI. (D) Mean hemifusion and fusion times versus number of fusogens (𝑛 = 40 simulations for each). (Mean waiting times computed assuming exponential distributions, see SI, Fig. S8.) Error bars: SEM. (E) Total pre-hemifusion membrane squeezing force versus numbers of fusogens (n = 40 simulations). Error bars: SD. (F) The entropy of a rod-like fusogen in a frozen environment is measured versus location of its LD C-termini. (G) Entropy for fixed-length LDs (tether) and worm-like chain LDs. Solid lines: rational function fits. (H) Time averaged thermal collision force on rod, and entropic force from fits in (G), versus rod location 𝑟. Rod force SDs: 5.1-10.8 pN (tether) and 4.4-14.1 pN (WLC).

Thus, molecular complexes that draw membranes together but lack rod domains aggregate between the vesicles and fail to drive membrane fusion.

### Rod-like fusogens generate forces that clear the fusion site and drive fusion

To highlight the effects of rod shape, we ran simulations using simplified model fusogens whose only structural feature is their rod shape, Fig. 2A. The simple nine bead fusogen has similar dimensions to SNARE complexes (2 nm wide, 10 nm long) but by design lacks structural details.

In simulations with six rod-like fusogens, fluctuating forces acted on each fusogen due to thermal collisions with membranes and other fusogens. These forces maintained the fusion site clear by pushing the rods outwards into a disordered ring. The outward radial forces stretched the LDs, inducing LD tension that squeezed the vesicles together and created a flattened contact zone with the rods at the perimeter (Figs. 4B, S4A). This configuration was established within a few μs.

These rod-mediated forces drove hemifusion and fusion. The mean hemifusion time was 0.7 ms, when a stalk formed at the contact zone boundary (Fig. 4B). Almost instantly, the fusogens migrated to and surrounded the stalk, but typically the stalk disintegrated after ∼ 0.1 ms, restoring the pre-hemifused state. After one or more of these reversible stalk episodes, a stalk matured to fusion when, after a delay, the cluster of fusogens surrounding the stalk expanded it into a hemifusion diaphragm (HD) and induced a localized simple pore in the HD that rapidly became a mature fusion pore (Fig. 4B; see Movies S3, S4 for an example of the full pathway). The mean fusion time was 2.8 ms (Figs. 4D, S4).

The time-averaged radial force that drove fusion was ∼ 6 pN per fusogen, Fig. 4C. In simulations with fusogens constrained in a small cluster near the vesicle-vesicle contact point, the mean force increased to ∼ 11 pN (Figs. 4C, S5). A requirement for these substantial forces is the rod-like structure, as globular fusogens generated only ∼ 3-4 pN outward forces whether constrained or not (Fig. 4C).

With more fusogens, forces were greater and fusion was faster (Fig. 4D, E). Three fusogens occasionally nucleated hemifusion stalks that always disintegrated. Five fusogens were required for fusion, and seven were sufficient to drive fusion in about a ms (Fig. 4D, S4). As the number increased from three to nine, the net pre-hemifusion squeezing force increased ∼ 3-fold to ∼ 140 pN (Fig. 4E). Whenever irreversible hemifusion occurred, the squeezing pressure of 9-11 atm (Fig. S4C) was similar to that required to irreversibly hemifuse DMPC bilayers on soft supports (70).

Thus, thermal collisions push rod-shaped fusogens outwards and squeeze the membranes. More rod-like fusogens produce a greater net squeezing force, giving faster hemifusion and fusion.

### The forces generated by rod-like fusogens are entropic

We next demonstrated that the origin of the forces that push rod-shaped fusogens outwards is the increased entropy in an expanded ring (47–49): a rod that moves radially outwards gains orientational entropy in the polar direction due to membrane curvature, and in the azimuthal direction due to increased separation from other rods (Fig. 4F).

To explicitly show the forces are entropic, we froze the system in a typical configuration, leaving the dynamics of one rod as normal (Fig. 4F). Discretizing the rod state space into 0.4 nm spatial cells and 12° angular cells, the rod configuration was recorded every ∼ 14 ns during ∼ 10 ms of simulation and the fraction of time spent in a given state was measured, 𝑃(𝑥, 𝑦, 𝑧, 𝜃, 𝜙), where (𝑥, 𝑦, 𝑧) locates the rod C-terminus and 𝜃 and 𝜙 are the rod orientational angles. The entropy 𝑆(𝑟) was computed as a function of the membrane anchoring location 𝑟 (see SI),

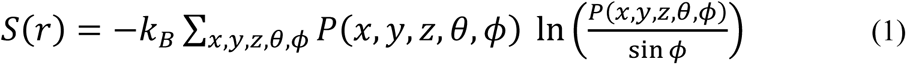

We first measured the entropy for fixed length floppy tether linkers. As expected, the entropy increased as the rod anchoring location moved outwards, Fig. 4G. We compared the entropic force to the measured force acting on the rod. Close to the membrane cleft, the entropic force 𝑓_ent_ = 𝑇 𝑑𝑆/𝑑𝑟 (𝑇 is temperature) accounted for most of the time averaged outward radial force on the rod, with a discrepancy we attribute to the soft tail of the rod-membrane and rod-rod potentials, Fig. 4H. With increasing distance, the rod force converged to the entropic force. Similar results were obtained for a worm-like chain LD spring law, Fig. 4G, H.

These results show that rod-like complexes are effective fusogens because rods generate high entropic forces that drive fusion. Rods are optimal entropic force generators due to their long reach and high excluded volume. By contrast, globular complexes generate small entropic force and become unproductively immobilized at the membrane-membrane contact point.

### Coarse-grained SNAREs fuse membranes via the same pathway as model rod-like fusogens

Next, we ran simulations using a systematically coarse-grained representation of the rod-like SNARE complex (47–49) (Fig. 2A). The complex has similar coarse dimensions to the model rod-like fusogens, but with SNARE-specific structural features.

Similarly to the model fusogens, entropic forces of ∼ 7-8 pN per SNARE complex maintained the fusion site clear and generated tensions of 21-22 pN in each LD, as we previously described (49). These tensions squeezed the vesicles together with a net squeezing force that increased from 60 to 170 pN as the number of SNAREs increased from 3 to 9 (Fig. 5C). As a result, the radius of the membrane contact zone, established within a few μs (Figs. 5A, S6), increased from ∼ 4 to ∼ 6 nm. The squeezing pressures of ∼ 13-16 atm catalyzed hemifusion by creating a stalk connection at the edge of the contact zone (Fig. 5A, C). Fusion was driven entropically by the SNAREs, which immediately surrounded the hemifused stalk connection and exerted even higher outward radial entropic forces (∼ 12 pN per SNARE for seven SNAREs) that expanded the stalk into a HD and opened a simple pore in the HD (Fig. 5A, D, Movie S5). Sometimes a reversible stalk preceded this final stalk-to-fusion episode (Fig. 5G).

**Figure 5.**
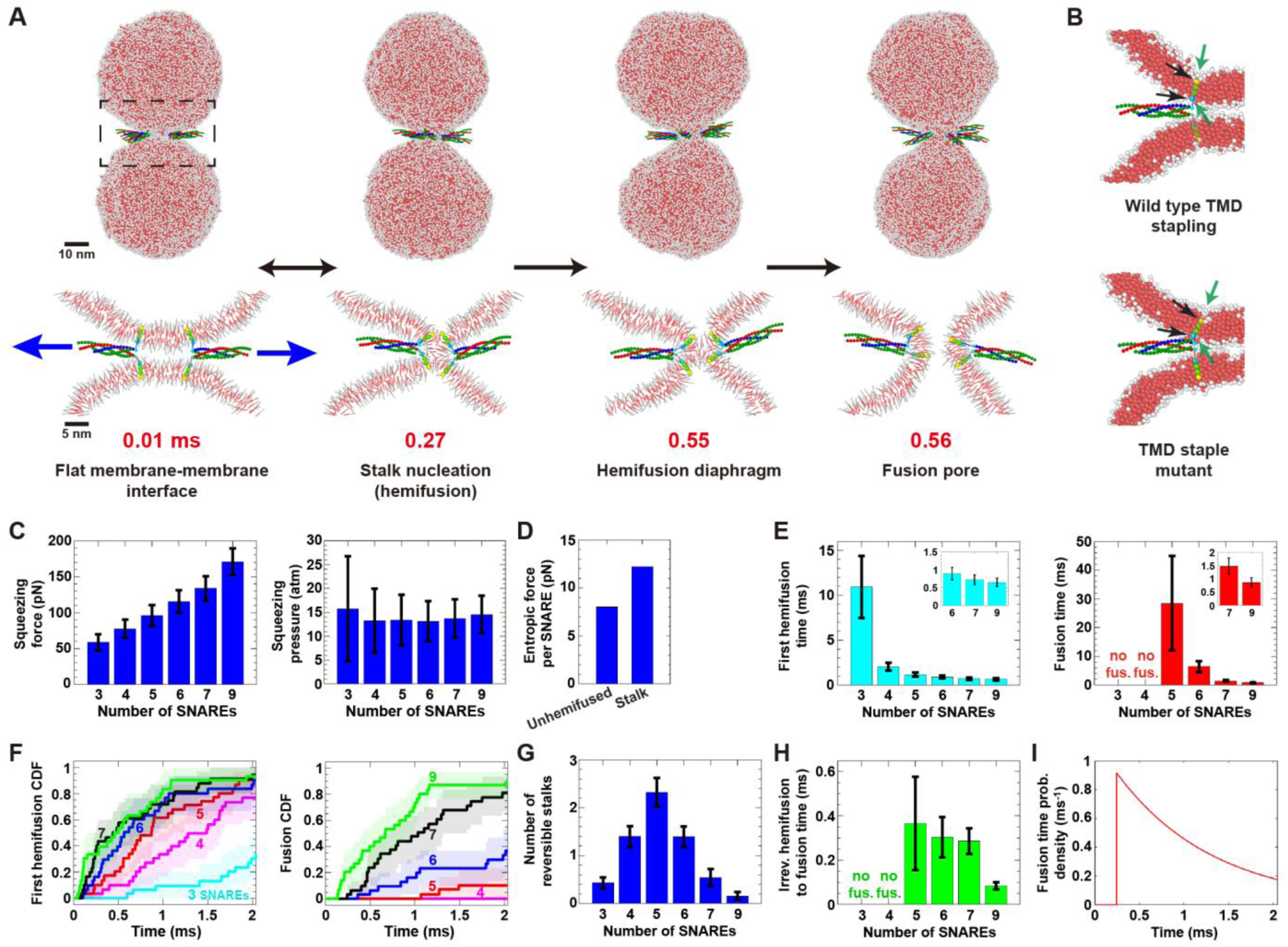
Coarse-grained SNARE fusogens generate entropic forces that drive membrane fusion along the same pathway as that for model rod-like fusogens. (A) Entropic forces (blue arrows) push seven SNAREs outwards into a ring, squeeze vesicles into a flat interface, nucleate a stalk at the interface edge, expand the stalk into a HD, and rupture the HD for fusion (see also Fig. S7A). (B) Top: Fusion is catalyzed by local membrane thinning (green arrows) by TMDs with staple beads (black arrows). Bottom: Thinning is abolished with mutants having neutral staples. No hemifusion or fusion occurred in ∼ 40 ms simulation. (C) Total pre-hemifusion squeezing force and pressure versus number of SNAREs (n = 10 simulations for each value). (D) Mean radial entropic force per SNARE during unhemifused and stalk phases (n = 10 simulations, seven SNAREs). (E) Mean first hemifusion and fusion times versus number of SNAREs. (F) Cumulative waiting time distributions for (E) (𝑛 = 30 simulations per distribution). (G) Mean number of reversible stalks during ∼ 2 ms of simulation versus number of SNAREs (n = 30 simulations per value). (H) Mean irreversible hemifusion-to-fusion time versus number of SNAREs (among ∼ 2ms simulations where fusion occurred). (I) Fusion time distribution for seven SNAREs (see Fig. S9). Error bars: SEM for (E), (H); SD for (C), (G). Shaded regions in (F): 95% confidence intervals. SDs in (D): 31.3 pN (unhemifused), 35.1 pN (stalk).

The greater entropic membrane squeezing forces generated by larger numbers of SNAREs yielded faster hemifusion and fusion (Figs. 5E, F, H, S7). Three or four SNAREs generated short-lived (< 0.1 ms) transient stalks, but no fusion in ∼ 60 ms of simulation. Five SNAREs were just sufficient for fusion: following several reversible stalk episodes (stalk lifetimes ∼ 0.3 ms), fusion occurred after a mean waiting time ∼ 30 ms (Fig. 5E, G). Nine SNAREs were enough to drive sub-ms fusion, with a mean fusion time of 0.9 ms either directly from the first stalk or, in ∼ 20% of cases, preceded by a single reversible stalk episode with lifetime ∼ 20 µs (Fig. 5E, G).

Thus, SNARE complexes and simplified rod-like fusogens fuse membranes along identical pathways, showing that the rod shape is the critical feature.

### TMD-induced membrane thinning is required to catalyze fast fusion

Reflecting the known structure of SNARE TMDs (15), the TMDs of simulated rod-like fusogens and SNAREs are somewhat shorter (∼ 3 nm) than the unperturbed membrane thickness (∼ 5 nm) and incorporate terminal cytosolic and luminal staple beads representing charged groups belonging to the TMD and juxtamembrane LD region that interact with the polar lipid head groups (59–62) (Fig. 2B). In simulations, this resulted in staple-enforced membrane thinning at TMD locations (Figs. 5B, S4D). At these sites of membrane dimpling the hydrophobic membrane interiors were partially exposed, raising the possibility that this effect could promote hemifusion and fusion.

To test if local membrane thinning promotes membrane fusion we switched the staples off, i.e. the attractive TMD staple-lipid head group interactions were abolished. This abolished membrane thinning (Figs. 5B, S4D). No hemifusion or fusion occurred in simulations with six rod-like fusogens or seven SNAREs (total running times ∼ 30 ms and ∼ 40 ms, respectively).

Thus, rod-mediated entropic forces drive fusion, but the catalytic effect of local TMD-mediated membrane disruption is indispensable for fast fusion.

### EFF-1 fusogens generate entropic forces that drive membrane fusion

Our results suggest the rod-like shape is the key property that makes SNARE complexes fusogenic. To further explore the possibility of a universal rod-mediated mechanism, we simulated a class II superfamily member, the eukaryotic cell-cell fusogen EFF-1, Fig. 2A. EFF-1 is rod-like, but in other respects its structure is quite different to that of SNAREs, having the characteristic class II 𝛽 sheet-rich structure and forming trimeric rods of similar length to SNARE complexes but much bulkier, Fig. 1. For each monomer, our model assumed the ten residues in the stem upstream of the TMD are unstructured and have the role of a LD.

A distinctive feature of EFF-1 is the acidic patch at the tip of the finger-like domain II of each monomer. In viral class II fusion proteins the *cd* loop connecting 𝛽 strands *c* and *d* is the hydrophobic fusion loop, whereas in EFF-1 the *cd* loop is negatively charged and, together with adjacent residues, constitutes the acidic patch thought to interact with polar lipid head groups (57, 58). It was proposed the patch serves to anchor the monomer upright in the membrane prior to trimerization (58).

Our model includes a coarse-grained version of the acidic patch, with characteristic tip-membrane interaction energy 𝜖_tip_ (Fig. 2A, Model section and SI). To determine this energy, we first simulated a single monomer anchored to a membrane, Fig. S10A, and found the value 𝜖_tip_ = 0.6 𝑘_𝐵_𝑇 best reproduced the experimentally measured orientational monomer fluctuations (58) (Fig. S10). With this value, seven EFF-1 trimers failed to drive hemifusion or fusion in ∼ 20 ms of simulation. Thus we ran simulations with variable 𝜖_tip_, revealing a catalytic effect that boosted fusion rates by local membrane disruption. Hemifusion and fusion times 𝜏 were exponentially sensitive, 𝜏 = 𝜏_0_𝑒^−𝜖tip/𝐸0^ with 𝜏_0_ ≈ 32 sec and 𝐸_0_ = 0.11 kT for hemifusion, and 𝜏_0_ ≈ 28 min and 𝐸_0_ = 0.09 kT for fusion, Fig. 6D. Extrapolating to the best fit value, 𝜖_tip_ = 0.6 𝑘_𝐵_𝑇, yielded EFF-1-mediated hemifusion and fusion times of 0.12 s and ∼ 2.5 s, respectively (Fig. 6D). It is interesting to note that EFF-1 fuses cells on min timescales during development (72).

**Figure 6.**
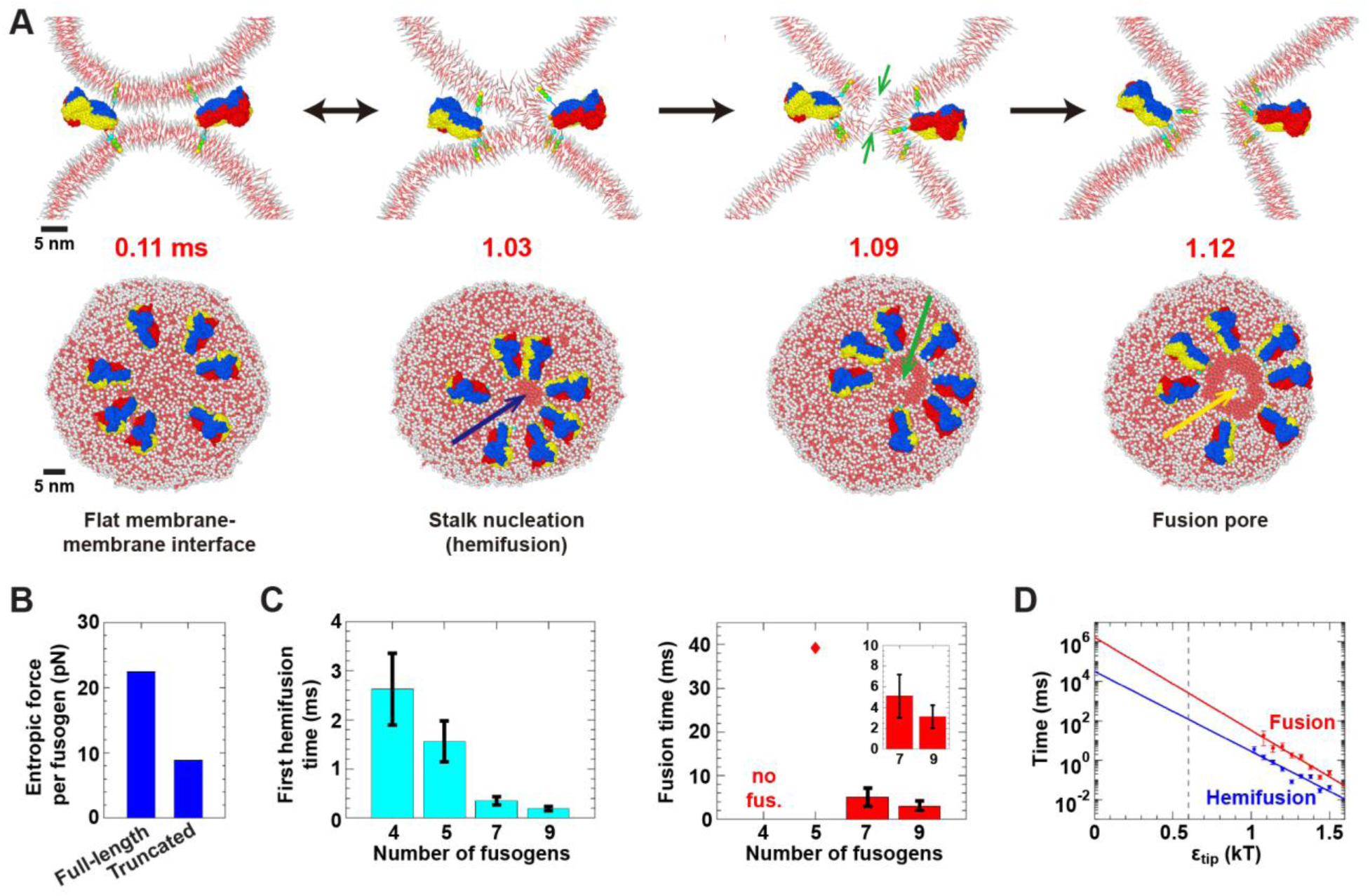
Coarse-grained EFF-1 fusogens drive fusion via a second entropic pathway. (A) Entropic forces maintain an expanded ring of seven fusogens, squeeze vesicles into a flat interface, nucleate a stalk (blue arrow) and elongate the stalk along the interface edge. Simple pores that open in each vesicle (green arrows) are encircled by the stalk, yielding a fusion pore (1.12 ms, yellow arrow). (B) Mean entropic force per EFF-1 with seven full-length or truncated EFF-1 complexes (𝑛 = 20 and 10 simulations, respectively; SDs: 66.5 pN and 58.7 pN). (C) Mean hemifusion and fusion times versus number of fusogens (𝑛 = 20 simulations per value). Diamond indicates a single event. (D) Hemifusion and fusion times versus acidic tip attraction 𝜖_tip_ (𝑛 = 20 simulations per value, seven fusogens). Solid lines: fits 𝜏 = 𝜏_0_𝑒^−𝜖_tip_/𝐸_0_^ with 𝜏_0_ ≈ 32 sec, 𝐸_0_ = 0.11 kT for hemifusion (𝑅^2^ = 0.89) and 𝜏_0_ ≈ 28 min, 𝐸_0_ = 0.09 kT for fusion (𝑅^2^ = 0.93). Dashed line: 𝜖_tip_ = 0.6 𝑘_𝐵_𝑇, value best reproducing experimental (58) EFF-1 monomer orientational distribution (see Fig. S10). Error bars: SEM for (C) and (D). For (A)-(C), 𝜖_tip_ = 1.2 kT.

The fusion pathway was similar to that for SNAREs and model rod-like fusogens. Seven trimers with 𝜖_tip_ = 1.2 𝑘_𝐵_𝑇 drove hemifusion and fusion after ∼ 0.35 ms and ∼ 5 ms, respectively. Large entropic forces of ∼ 22 pN per fusogen pushed the fusogens into an expanded ring, squeezed the vesicles and nucleated a stalk at the boundary of the membrane contact zone (Fig. 6A, B). Following one or more reversible stalk episodes, a stalk was elongated along the boundary by fusogens lining the stalk edge (Fig. 6A). Two simple pores developed near the stalk, one in each vesicle, and fusion was completed when the stalk encircled the pores to make a fusion pore (Fig. 6A, Movie S6). Hypothetical non-rod-like EFF-1 mutants truncated to ∼ 2.3 nm length generated lower entropic forces of ∼ 9 pN per fusogen, accumulated between the vesicles, and failed to hemifuse or fuse them (Figs. 6B, S11, Movie S7).

In conclusion, despite major structural differences, EFF-1 generates entropic forces that drive fusion similarly to SNARE complexes. Again, the coarse rod shape is the key structural feature for fusogenicity. EFF-1 uses the same pathway, except stalks expand longitudinally rather than isotropically (cf. Figs. 5A, 6A). Fusion is accelerated by local membrane disruption by the charged EFF-1 trimer tip, analogously to local disruption by TMD-mediated membrane thinning for SNARE-mediated fusion.

## Discussion

### A universal fusion mechanism has been a driver of structural convergence among fusogens

In the final state that drives fusion, numerous eukaryotic and viral fusogens are rod-like *trans*-complexes (Fig. 1). Our simulations suggest the rod shape is the essential mechanistic feature, because rods generate high entropic forces that drive hemifusion and fusion (Fig. 7). Rods are optimally efficient entropic force generators due to their long reach, which endows them with high excluded volume: for complexes of length 𝐿 and width 𝐷 of a given complex volume ∼ 𝐿𝐷^2^, the mutual excluded volume ∼𝐿^2^𝐷 is greater if the length is greater (Fig. 7).

**Figure 7.**
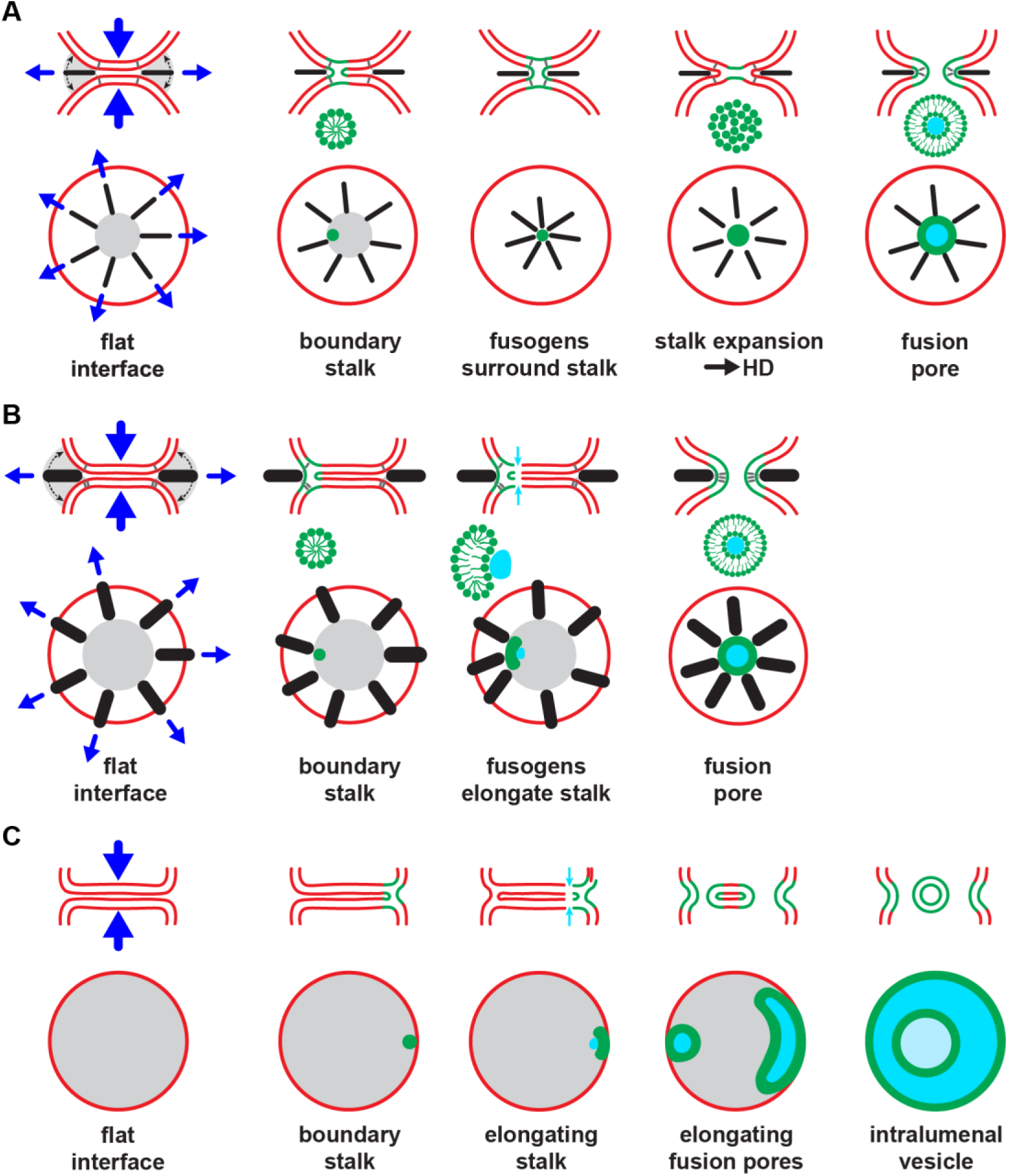
Three membrane fusion pathways. (A) SNAREs and class I superfamily fusogens, slender α-helical rods. Our simulations suggest these fusogens use a common pathway to fuse small membrane compartments. Entropic forces (arrows) squeeze membranes and drive a boundary hemifusion stalk. The rods then surround and entropically expand the stalk into a fusion pore. (B) Class II fusogens, bulky β-sheet-rich rods. This pathway is similar to (A), but following hemifusion the bulky fusogens cannot surround the stalk and instead line its edge and entropically expand it longitudinally. Simple pores rapidly form in each vesicle that the stalk encircles to make a fusion pore. (C) Brute force. A third pathway may be used when large interfaces are created, e.g. SNARE-mediated fusion of vacuoles in plants and fungi. The pathway is similar to (B), but several fusion pores elongate along the boundary edge and merge to release an intralumenal vesicle.

Despite differences in structural detail, coarse-grained SNARE complexes, EFF-1 fusogens, and model rod-like complexes generated entropic forces that drove fusion along identical or similar pathways (Figs. 4B, 5A, 6A). The entropic forces relied on the coarse rod-like shape of the fusogens. Drawing the membranes together was insufficient: globular fusion complexes or truncated versions of rod-like fusogens achieved membrane proximity but failed to generate substantial entropic forces and failed as fusogens (Figs. 4A, C, 6B, S11).

Based on these results, we suggest the structural similarity between SNAREs and class I superfamily fusogens is an instance of convergent evolution, where independent evolutionary routes arrived at the same solution to the fusion challenge, namely rod-generated entropic forces. Alternatively, the origin could be the transfer of a structural fusion fold between host and virus (73). However, arguing against this is the fact that the structural similarity is only that both ectodomains have 𝛼-helical rod domains: SNARE complexes are coiled-coils of four 𝛼-helices (14, 15), whereas class I fusogens have coiled-coil backbones of three 𝛼-helices, part of which is complemented by three anti-parallel 𝛼-helices in a six-helix bundle (6, 17). We propose that the origin is the high entropic force that rod-like structures can generate for fusion.

Generally, eukaryotic and viral fusion proteins have not evolved independently, as numerous transmissions have occurred between eukaryotic hosts and viruses (19, 74). In class I, sequence analysis showed that syncytins in different species resulted from distinct insertions from different viruses (19, 75). In class II, the oldest known fusogen is HAP2. It was suggested HAP2 could be the ancestral eukaryotic fusogen from which class II viral fusogens evolved, or alternatively HAP2 may have begun as a viral fusogen that later integrated into eukaryotic genomes and evolved into the gamete fusogen (74). The class II eukaryotic fusogen EFF-1 we simulated here is a trimer assembled from identical monomers anchored in opposing membranes and interacting in *trans*. Since trimers would not naturally be expected in the symmetric situation of a bilaterally acting fusogen, this suggests a viral origin (57). While many class II fusion proteins belong to members of closely related virus families (76), others have been discovered in unrelated families (73, 77). Moreover, in some cases different viruses within the same family have different fusion proteins, some in class II and some not (78), suggesting that some viruses have independently obtained their class II fusion proteins from different eukaryotic hosts, or from other viruses during coinfections (73).

Thus, fusogens with rod-like structure have repeatedly been transferred in multiple apparently independent events, suggesting a universal rod-based fusion mechanism may have been the evolutionary driver. More broadly, the mechanism may have driven convergence toward rod-like structure among large classes of fusogens.

### SNAREs and class I fusogens may use a common fusion pathway

In simulations, SNARE complexes and simple rod-shaped fusogens followed the same pathway to fuse small vesicles (Figs. 4B, 5A). The common feature selecting the pathway was that both are long slender rods, independent of structural details (Fig. 2A). This suggests SNARE complexes and class I superfamily fusogens use the same pathway to fuse a given pair of membrane compartments, since all are long 𝛼-helical rods (Fig. 7A).

In this pathway (49), rod-generated entropic forces push the fusogens outwards so the LDs become stretched with high tensions (∼ 21 pN for SNARE complexes) that squeeze the vesicle membranes into a flat membrane-membrane interface with the fusogens at the boundary, Fig 7A. This quasi-equilibrium situation typically persists for some fraction of a ms, until the squeezing forces induce a stalk at the interface boundary that is stabilized by one or more fusogens that locally squeeze the membranes, reducing the stalk height closer to its energetically preferred value. Within 𝜇s of stalk nucleation, the fusogens regroup around the stalk and, being densely clustered, exert even greater outward entropic force, expanding the stalk into a HD and rupturing the HD for fusion (Fig. 7A, Movie S5). Sometimes this sequence is punctuated by a disassembling stalk (hemifission, reversible stalk) (Fig. 5G).

Consistent with the intermediates on this pathway, large bilayer interfaces and HDs were visualized between liposome pairs in reconstituted SNARE systems (39, 40). Hemifusion lies on the fusion pathway in pneumocytes (44), while HDs were observed between the PM and synaptic vesicles, and between the PM and secretory granules in chromaffin and pancreatic β cells (∼ 200 nm) (41, 43). Mediated by the class I fusogen haemagglutinin (79), HDs were visualized between influenza virus-like particles and liposomes.

### A second entropic fusion pathway used by class II fusogens

Like SNARE complexes and class I fusogens, class II fusogens are rod-like. However, the rods are far bulkier (Fig. 1). In simulations, the eukaryotic class II EFF-1 fusogen generated entropic forces that drove fusion similarly to SNAREs, but following a slightly different pathway because EFF-1 is too bulky to fit between the membranes and surround the stalk, instead being confined to its outside edge. As a result, the stalk at the interface boundary is elongated by fusogens lining its edge, Fig. 7B, in contrast to the SNARE/class I pathway where fusogens expand the stalk isotropically, Fig. 7A. Fusogen-induced strains then triggered two small pores which were then encircled by the elongating stalk to form a fusion pore.

Our results suggest all class II fusogens use this pathway, as the bulky rod shape is the determinant. Detailed pathways are not experimentally demonstrated, but hemifusion appears to be an intermediate, as stalk intermediates were reported in EFF-1-mediated vesicle-vesicle fusion (58), and HDs were visualized between liposomes and chikungunya virions, mediated by class II fusogens (80).

### Fusion by brute force uses a third fusion pathway similar to the vacuole fusion pathway

In simulations where two vesicles were simply pressed together, the interface was much larger than with fusogens and all hemifusion and fusion processes occurred at the high curvature interface boundary, Fig. 7C. High enough forces nucleated a stalk at the boundary, which grew longitudinally and then encircled two pores that formed, one in each vesicle. The resulting fusion pore elongated along the boundary and combined with other similar pores, cutting free an intralumenal vesicle. At intermediate stages, partial cutting generated intralumenal tubes or even networks.

This pathway has compelling parallels with that for SNARE-driven vacuole fusion. Vacuoles are micron-sized membrane-bound organelles widely studied in yeast, plants and other organisms (71, 81). Docked vacuoles form large, flattened membrane-membrane interfaces, with docking and fusion machinery along the interface boundary (82). A hallmark of vacuolar fusion is the formation of an intralumenal vesicle within the fused vacuole, whose function may be to maintain vacuole shape by regulating its area-to-volume ratio (42, 82). Experiments suggest a mechanism whereby the vesicle is released intralumenally by hemifusion and fusion pore opening at the boundary followed by pore expansion along the boundary (42, 71, 82). This is very similar to the brute force-mediated vesicle release mechanism (Figs. 3, S2). We propose this pathway is used by SNAREs and other slender rod-like fusogens to fuse micron-sized compartments with extended preformed interfaces.

### Parallels with calcium-driven fusion

In the SNARE/class I fusion pathway, fusogens adhere two membranes in an extended interface and subsequently surround and entropically expand the stalk into a HD and rupture the HD for fusion (Fig. 7A). This has parallels with the pathway driven by Ca^2+^, which has long served as a model protein-free system for biological fusion (83, 84). Divalent cations such as Ca^2+^ adhere negatively charged GUVs to other GUVs (83) or suspended bilayers (84). Analogously to entropic forces, membrane tension and selective contraction of the outer leaflets that contact Ca^2+^drive HD expansion and rupture the HD (85–87).

The entropic and electrostatic driving forces are quite different. Nonetheless, both pathways involve a membrane-adhered state, a growing HD phase and HD rupture. Further, during Ca^2+^-driven fusion if the HD stochastically fails to rupture during its fast expansion phase, it continues growing and equilibrates to a dead-end state. The equilibrium size distribution of these large dead-end HDs was measured (83) and shown to be related to the driving forces (85–87). Similarly, very large HDs between vacuoles (42) and between liposomes in reconstituted SNARE systems (39, 40) may be dead-end or slow secondary pathways.

### Local membrane-disrupting effects catalyze membrane fusion

While entropic force was the driver of fusion, local membrane-disrupting effects played an equally vital catalytic role by lowering fusion energy barriers. During simulated SNARE- or model rod-like fusogen-mediated fusion, TMD-mediated membrane thinning (88, 89) at the interface boundary dimpled bilayers and, together with the high curvature at the boundary due to entropic squeezing, exposed hydrophobic lipid tails and accelerated hemifusion at least ∼ 50-fold (Figs. 5B, S4D). This is in accord with a Martini study of isolated SNARE TMDs suggesting TMD-mediated membrane thinning may be associated with lowering of the stalk activation energy (89). More broadly, mutations of VAMP SNARE TMDs reduced or abolished fusion (90, 91). Analogously, fusion peptides and loops in viral fusogens likely promote fusion not only by bridging membranes, but also by modulating bilayer properties (92).

Similarly, EFF-1 drove fusion entropically, but local interactions of the negatively charged EFF-1 tip with the membrane (57, 58) strongly accelerated fusion, Fig. 6D. AFF-1 is a structurally homologous class II eukaryotic cell-cell fusogen lacking the charged tip; extrapolating our fit, Fig. 6D, to zero tip-membrane interaction energy suggests a ∼ 30 min AFF-1-mediated fusion time. Overall, regulation of fusogen tip strength may allow modulation of fusion times from ms to min.

### Fusion is a waiting game of ms or longer

We find membrane fusion is microscopically slow: on 𝜇s timescales, the SNARE complexes and membranes attain a quasi-equilibrated state, long before fusion which requires ms timescales. This dynamic metastable state is maintained for about a ms, when continuously fluctuating entropic forces impose a certain probability per unit time to hemifuse and fuse; fusion is probabilistic, with a broad roughly exponential waiting time distribution with mean of order a ms (see Figs. 5I, S9).

To our knowledge, this is consistent with all published electrophysiological measurements at neuronal synapses, where broad distributions of vesicle release times are reported with widths of order a ms or longer (29, 30). As for any broad distribution, there are some very early events, but these are atypical; thus, the delay time (first post-synaptic response) is not representative of the release time (31). This extensive data argues against alternative proposed mechanisms where fusion occurs almost immediately following SNARE complex formation, which likely occurs on 𝜇s timescales once the participating 𝛼-helices are released (28).

Being slow, physiological fusion is inaccessible to atomistic MD simulations or mildly coarse-grained approaches. These methods are critical to study local interactions and small timescale effects, but to capture the collective molecular behavior that yields membrane fusion on ms timescales requires MD methods that use radical coarse-graining such as in the present study.

### More fusogens produce higher entropic forces and drive faster fusion

A common view has been that fusion requires a certain number of SNARE complexes and occurs rapidly if the requirement is satisfied (32–35). In our simulations, a different phenomenology emerged: regardless of the fusogen - SNARE complexes, EFF-1 trimers or simple rod-like fusogens - more fusogens generated higher entropic forces and faster hemifusion and fusion (Figs. 4D, E, 5C, E, 6C). This behavior is a corollary of fusion being a long waiting process rather than “all-or-nothing,” since the long wait is characterized by a fusion probability per unit time, which inevitably depends on how many fusogens are present.

Faster fusion with more fusogens has interesting implications for synaptic transmission and plasticity. In neurons, transient action potential-evoked calcium influx at presynaptic terminals unclamps SNARE complexes to fuse synaptic vesicles in the readily releasable pool to the plasma membrane (3). We thus expect release rates and the net vesicle release probability 𝑃_ves_ during the calcium transient to depend on the number of unclamped SNAREs. Indeed, such a dependence has been observed in electrophysiological studies with decreased (51) or increased (50) numbers of active SNARE complexes mediating neurotransmitter release, when 𝑃_ves_ and release rates were decreased or increased, respectively. More generally, our work raises the possibility that regulation of the number of active SNAREs and their fusogenicities could serve as a means to modulate synaptic strength.

## Acknowledgements

Research reported in this publication was supported by the National Institute of General Medical Sciences of the National Institutes of Health under award number R01GM117046 to B.O’S. The content is solely the responsibility of the authors and does not necessarily represent the official views of the National Institutes of Health. We acknowledge computing resources from Columbia University’s Shared Research Computing Facility project.

## Author contributions

B.O’S designed the research. I.C.B., J.Z, and D.A performed the simulations. B.O’S., I.C.B., J.Z., D.A., Z.M and K.D. analyzed the data. B.O’S., I.C.B., J.Z., D.A., and Z.M interpreted the data and wrote the paper.

## Data availability

Simulation results supporting the findings of this paper, and codes to perform the simulations, to analyze the data, and to generate the technical figures are available in the Zenodo repository (DOI: 10.5281/zenodo.15366357). Because of their large size, simulation trajectory files are not included in the repository but are available upon request from the corresponding author. The codes are also available in the GitHub repository (https://github.com/OShaughnessyGroup-Columbia-University/fusogens_rod_shaped).

