## Supporting Information for "Why are so many fusogens rod-shaped?"

### Supporting Information Text

#### EFF-1 coarse-graining procedure

Coarse-grained EFF-1 fusogens were designed to approximate the solvent excluded surface of the post-fusion EFF-1 complex (PDB: 4OJC) (1). Using PyMOL (2), we generated a surface mesh with a solvent radius of 0.44 nm and applied a negative offset of 0.44 nm to it. Beads of 0.44 nm radius were placed on the mesh vertices using a random sequential adsorption (RSA) algorithm, enforcing a minimum separation of 0.44 nm between bead centers. The process continued until no further beads could be added, yielding a total of 630 beads. For visualization, beads were sorted and colored by protomer, with each bead assigned to the protomer containing the closest residue center of mass.

Ten residues upstream of the EFF-1 transmembrane domains (TMDs) were assumed unstructured prior to fusion and modeled as linker domains (LDs). The EFF-1 LDs thus consist of residues I552-D561, representing the last ten residues of the stem region following β strand *n* (1). The LDs of our coarse-grained EFF-1 fusogens extend from the center of mass of the first LD residue (I552) in each protomer to the center of the cytosolic TMD staple.

In EFF-1, the loop corresponding to the hydrophobic fusion loop of viral class II fusogens contains a negatively charged segment (178-SEDD-181) facing the membrane. Together with D136 located nearby, these residues form an acidic patch thought to interact with polar lipid head groups (1, 3). Our model includes a coarse-grained representation of this acidic tip, consisting of one bead per protomer positioned at the center of mass of the residues comprising the acidic patch (Fig. 2A, orange beads). These beads interact attractively with lipid head groups and have no other interactions (see below).

#### Non-bonded interactions

With three exceptions listed below, all beads in the simulation interact repulsively through the Weeks-Chandler-Andersen (WCA) potential, as described in (4) and in our previous study (5).

Exceptions: (1) Solvent beads inside vesicles (see below) do not repel one another. (2) The SNARE LDs connect the TMDs to two ghost beads located at the C termini of the VAMP and syntaxin SNARE motifs (Fig. 2A, cyan beads). These ghost beads have the same diameter as other SNARE beads but do not participate in any other interactions. Similarly, the LDs of the model rod-like fusogen connect the TMDs to two ghost beads at the C terminus of the rod domain (Fig. 2A, cyan beads), which have the same diameter as the TMD beads. (3) The beads representing the acidic patch at the tip of EFF-1 (Fig. 2A, orange beads) have attractive interactions with lipid head beads (see below) and no repulsive interactions.

Additional attractive interactions are included among lipid tail beads (4), between lipid tail beads and TMD hydrophobic beads, among hydrophobic TMD beads, and between the two hydrophilic TMD staple beads and lipid head beads, as described in our previous study (5).

The three beads representing the acidic patch at the tip of EFF-1 interact attractively with lipid head beads, using the same potential form as the other attractive interactions in the simulation:

$$V_{\mathrm{tip}}(r) = \begin{cases} -\epsilon_{\mathrm{tip}}, & r < r_{\mathrm{c}} \\ -\epsilon_{\mathrm{tip}} \cos^2 \dfrac{\pi(r - r_{\mathrm{c}})}{2w_{\mathrm{c}}}, & r_c \le r \le r_{\mathrm{c}} + w_{\mathrm{c}} \\ 0, & r > r_{\mathrm{c}} + w_{\mathrm{c}} \end{cases}$$

where $r_c = 2^{1/6}(D_{\text{head}} + D_{\text{EFF-1}})/2$ (see Table S1 for bead diameters $D_{\text{head}}$ and $D_{\text{EFF-1}}$), $w_{\text{c}} = 1.6(D_{\text{head}} + D_{\text{EFF-1}})/2$, and the attractive well depth $\epsilon_{\text{tip}}$ was varied in simulations (see below).

**Bonded interactions**

The bonds between the four beads making up each lipid followed (6) and were described in our earlier study (5).

For the model rod-like fusogen, each linker domain connects the center of the cytosolic TMD staple bead to the center of a ghost bead at the rod's C terminus (Fig. 2A, cyan beads). For non-rod-like fusogens, the LDs connect the center of the cytosolic TMD staple directly to the center of the globular domain. The LDs of the model rod-like and non-rod-like fusogens are represented by fixed-force springs with a tension of 18 pN down to 0.1 nm extension, with the force decreasing linearly to zero below this threshold.

For coarse-grained SNAREs, the LDs extends from the surface of hydrophobic TMD bead adjacent to the cytosolic staple to the center of a ghost bead at the C terminus of the Stx or VAMP SNARE motifs (Fig. 2A, cyan beads). The unstructured LDs of SNAREs and EFF-1 are represented by identical worm-like chains, using parameters previously tuned for SNAREs in our previous study (5).

**Simulation procedures**

All simulations feature two 50 nm diameter synaptic vesicles in a 88 nm x 88 nm x 123 nm box with periodic boundary conditions. Within vesicles a fictitious gas of ghost beads creates pressure $P_{\text{ves}}$ set by bead density according to the ideal gas law. We used the pressure to regulate the tension of the vesicle of radius $R_{\text{ves}}$ according to the Young-Laplace law, $\gamma_{\text{ves}} = P_{\text{ves}} R_{\text{ves}}/2$, with $\gamma_{\text{ves}} = 0.05$ pN/nm in simulations with fusogens and 1 pN/nm in brute force simulations.

For simulations using brute force to squeeze the vesicle membranes together, we placed two equilibrated vesicles within a cylinder extending along the z-axis, with radius $R_{\text{cyl}} = 30$ nm (Fig. S1). The cylindrical wall interacts with lipid beads via a repulsive potential:

$$V_{\text{cyl}}(r) = \begin{cases} 4\epsilon_{\text{wall}} \left(\dfrac{\sigma_{\text{wall}}}{r}\right)^{12}, & r < r_{\text{c}} \\ 0, & r \geq r_{\text{c}} \end{cases}$$

where $\epsilon_{\text{wall}} = 0.06\ k_B T$ and $\sigma_{\text{wall}} = 0.44$ nm, equal to the radius of a lipid tail bead, and $r_{\text{c}} = 0.88$ nm. To generate the initial condition, a snapshot of an equilibrated vesicle was obtained (Fig. S1), and two copies of this vesicle were placed such that their outer leaflets were initially 4 nm apart along the z-axis. A constant force of magnitude $f_{\text{const}}$ was then applied to every lipid bead to press the two vesicles together throughout the ~ 1.4 ms simulation time.

All simulations with rod-like fusogens, non-rod-like fusogens, SNARE complexes, and EFF-1 fusogens were started from an initial setup with two ~ 50 nm diameter vesicles with crystalized bilayers and the fusogens placed evenly in a ring of ~ 7 nm inner radius around the point of closest approach between the vesicles (Fig. S6). The system was equilibrated for 1 μs with a time step of 1/100$^{\text{th}}$ of that in the actual simulation. Simulations with model rod-like and non-rod-like fusogens were then run for ~ 1.4 ms. Simulations with SNAREs and EFF-1 fusogens were run for ~ 2 ms.

In a subset of simulations featuring rod-like or non-rod-like fusogens, we fixed the positions of the TMDs to impose a fixed-radius fusogen cluster, leaving the dynamics of all other components as normal (see Fig. S5).

Simulation trajectories were visualized with the Open Visualization Tool (OVITO) (7).

**Membrane squeezing force and pressure measurements**

The total membrane squeezing force was calculated by summing the vertical components of the LD zippering forces from each fusogen, which are roughly perpendicular to the membrane-membrane interface. Measurements were conducted every 200,000 steps for rod-like fusogens and every 200 steps for SNAREs and EFF-1 fusogens.

The radius of the flat membrane-membrane interface was defined as the radius of the fusogen ring. For rod-like fusogens, this radius was calculated as the average distance between the x-y projections of the LD N-termini (where the LDs connect to the rod) and their centroid. Similarly, for SNARE complexes, the radius was measured as the average distance between the x-y projections of the LD C-termini (where the LDs connect to the TMDs) and their centroid. The ring radius was measured every 200,000 steps for rod-like fusogens and every 200 steps for SNAREs.

The membrane squeezing pressure was computed every 200,000 steps for rod-like fusogens and every 200 steps for SNAREs, using the corresponding total membrane squeezing force and ring radius at each time point. Values were then averaged across all time points up to the first hemifusion event and across multiple simulations (n = 40 for rod-like fusogens, n = 10 for SNAREs).

**Entropic force measurements**

The net force acting on each fusogen and the LD stretching forces were measured every 200 steps. The force generated by excluded volume interactions among fusogens and between fusogens and membranes is the vector difference between the net force acting on the fusogen and the LD stretching forces, and the entropic force is the radial component of this force.

The radial unit vector for each rod-like fusogen points from the center of the fusogen ring (defined in the previous section) to the midpoint of the x-y projections of the two LD N-termini. For the globular fusogen, the radial unit vector points from the center of the fusogen ring to the x-y projection of the globular domain's center. For SNAREs the radial unit vector points from the center of the fusogen ring to the midpoint of the x-y projections of the two LD C-termini. For EFF-1 the radial unit vector points from the center of the fusogen ring to the centroid of the x-y projections of the three LD C-termini.

Fig. 4C shows the outward entropic force per fusogen from simulations with six rod-like or globular fusogens, both under normal conditions and with the fusogen TMDs constrained in a ~ 3 nm ring at the fusion site (see Fig. S5). Since the entropic force is highly fluctuating due to its hard core nature, SDs are omitted from Fig. 4C for clarity. SDs for unconstrained and constrained simulations, respectively: 16.1 pN, 20.4 pN (rod-like fusogens), and 22.1 pN, 29.6 pN (globular fusogens).

**Entropy measurements**

The entropy of a model rod-like fusogen was measured in simulations where all components were frozen except for the rod domain of a single fusogen. This was done to ensure a constant configurational phase space for the rod. The frozen system configuration was obtained from a simulation with six rod-like fusogens with regular dynamics.

The C-termini of the LDs, where the LDs would connect to the TMDs were incrementally shifted radially outwards, away from the edge of the membrane interface, from $r$ = 0 nm to $r$ = 6 nm in steps of 0.2 nm, where $r$ = 0 nm corresponds to the native membrane anchoring location in the simulation frame used as the frozen configuration (Fig. 4F, left). The radial direction was defined as the average orientation of the fusogen LD force in the x-y plane during a simulation with $r$ = 0 nm.

At each position we conducted five simulations, each ~ 2 ms long. The rod's position $(x, y, z)$ and orientation $(\theta, \phi)$, as well as the net force acting on the rod due to rod-rod and rod-membrane interactions, were recorded every 200 steps (~ 14 ns) (n ≈ 750,000 measurements). The rod entropy was calculated as:

$$S(r) = -k_B \sum_{x,y,z,\theta,\phi} P(x, y, z, \theta, \phi) \ln\left(\frac{P(x, y, z, \theta, \phi)}{\sin\phi}\right).$$

Here $P(x, y, z, \theta, \phi)$ is the probability of observing a given rod configuration, with positions and orientations binned into cells of size 0.4 nm and 12°, respectively. The factor $\sin\phi$ accounts for the density of states, since the surface element on the unit sphere is $dA = d\theta\ d\phi \sin\phi$.

The following potential was used for the LDs:

$$V_{\mathrm{LD}}(r) = k_B T \frac{l_c}{l_p} \frac{\varepsilon_{\mathrm{LD}}}{4\left(1 - \frac{r}{l_c}\right)^m} \left(3\left(\frac{r}{l_c}\right)^2 - 2\left(\frac{r}{l_c}\right)^3\right)$$

where $l_c$ = 3.65 nm is the contour length, $l_p$ = 0.5 nm is the persistence length, and $\varepsilon_{\mathrm{LD}}$ and $m$ are adjustable parameters. Simulations were performed using two LD potentials: a nearly fixed-length tether ($\varepsilon_{\mathrm{LD}}$ = 5e-5, $m$ = 3) and a worm-like chain identical to that used in SNARE and EFF-1 simulations ($\varepsilon_{\mathrm{LD}}$ = 1, $m$ = 1).

**Estimation of mean times for events along the fusion pathway**

Since hemifusion and fusion did not occur in all runs, depending on the simulation conditions, mean waiting times were computed assuming exponential distributions. The survival probability curve was fit to $e^{-t/\tau}$, where $\tau$ is the mean time (Fig. S8).

This fitting procedure was repeated for 200 bootstrap resamplings, each generated by sampling $n$ simulations with replacement from the total $n$ simulations. The reported mean time $\langle\tau\rangle$ is the average of the fitted $\tau$ values across these 200 resamplings. The reported SEM is calculated as $\langle\tau\rangle/\sqrt{N_{\mathrm{obs}}}$, where $N_{\mathrm{obs}}$ is the number of simulations in which the event of interest (first hemifusion, irreversible hemifusion, or fusion) was observed.

With 5 EFF-1 fusogens, fusion was observed in only 1/20 simulations. In this case, the mean fusion time was calculated as $t_{\mathrm{avg}} = t_{\mathrm{obs}} + (N_{\mathrm{sim}} - 1)t_{\mathrm{sim}}$, where $t_{\mathrm{obs}}$ is the observed waiting time for fusion in the successful simulation, $N_{\mathrm{sim}} = 20$ is the total number of simulations, and $t_{\mathrm{sim}} \approx$ 2 ms is the duration of each simulation.

**Simulations of EFF-1 monomers**

It has been proposed that the acidic patch at the tip of EFF-1 anchors monomers in an upright configuration on the membrane prior to trimerization (3). In our simulations, we tuned the strength of the attractive interaction between beads representing the EFF-1 acidic tip and lipid head beads to reproduce the experimentally measured orientational distribution of membrane-anchored EFF-1 monomers reported in (3).

We generated a coarse grained representation of a single EFF-1 monomer based on the crystal structure (PDB: 4OJC) (1), using the same approach described for the trimer to approximate the solvent-excluded surface of the monomer with 0.44 nm radius beads. A bead representing the acidic tip was included at the center of mass of the acidic patch residues, as described above for the trimer. Unlike in the trimer model, the stem of the monomer was assumed structured up to the last resolved residue in the crystal structure, T560, located one residue upstream of the TMD (1).

The monomer was tethered to a TMD identical to those of the trimer via a fixed-force spring exerting a tension $f_{\mathrm{LD}}$, which extended from the center of mass of T560 to the center of the cytosolic TMD staple. We simulated a single EFF-1 monomer with its TMD embedded in a planar membrane under a tension $\gamma_{\mathrm{PM}} = 0.5$ pN/nm, and recorded its orientation relative to the membrane every 200 steps (~ 7 ns) (Fig. S10A).

To reproduce the experimental orientational distribution, we performed a parameter scan varying both the attractive well depth for the tip-lipid head bead interaction, $\epsilon_{\mathrm{tip}}$ $(0 - 1.8\ k_B T)$, and $f_{\mathrm{LD}}$ (5-25 pN), with two simulations of ~ 1 ms each per parameter pair. The upright orientation was defined to visually match that reported in (3) (Fig. S10B). For each parameter pair, we calculated: (i) the mean tilt offset, defined as the tilt of the average monomer orientation relative to the upright reference (Fig. S10D), and (ii) the tilt spread, defined as the mean deviation of the monomer from its average orientation (Fig. S10E). In ref. (3), the mean orientation was upright (zero mean tilt offset), and the tilt was isotropic with a mean magnitude of 33±17° relative to the membrane normal. These experimental values were best reproduced in simulations with $\epsilon_{\mathrm{tip}} = 0.6\ k_B T$ and $f_{\mathrm{LD}} = 10$ pN (Fig. S10C-F).

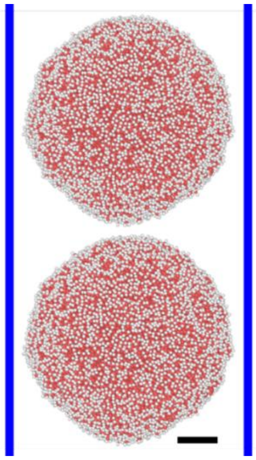

**Figure S1.** Side view of the simulation initial condition for fusion via brute force. Two ~ 50 nm diameter vesicles are confined within a cylinder (blue lines) to prevent them from sliding past each other when pressed together by a constant force applied to every lipid bead. Scale bar: 10 nm.

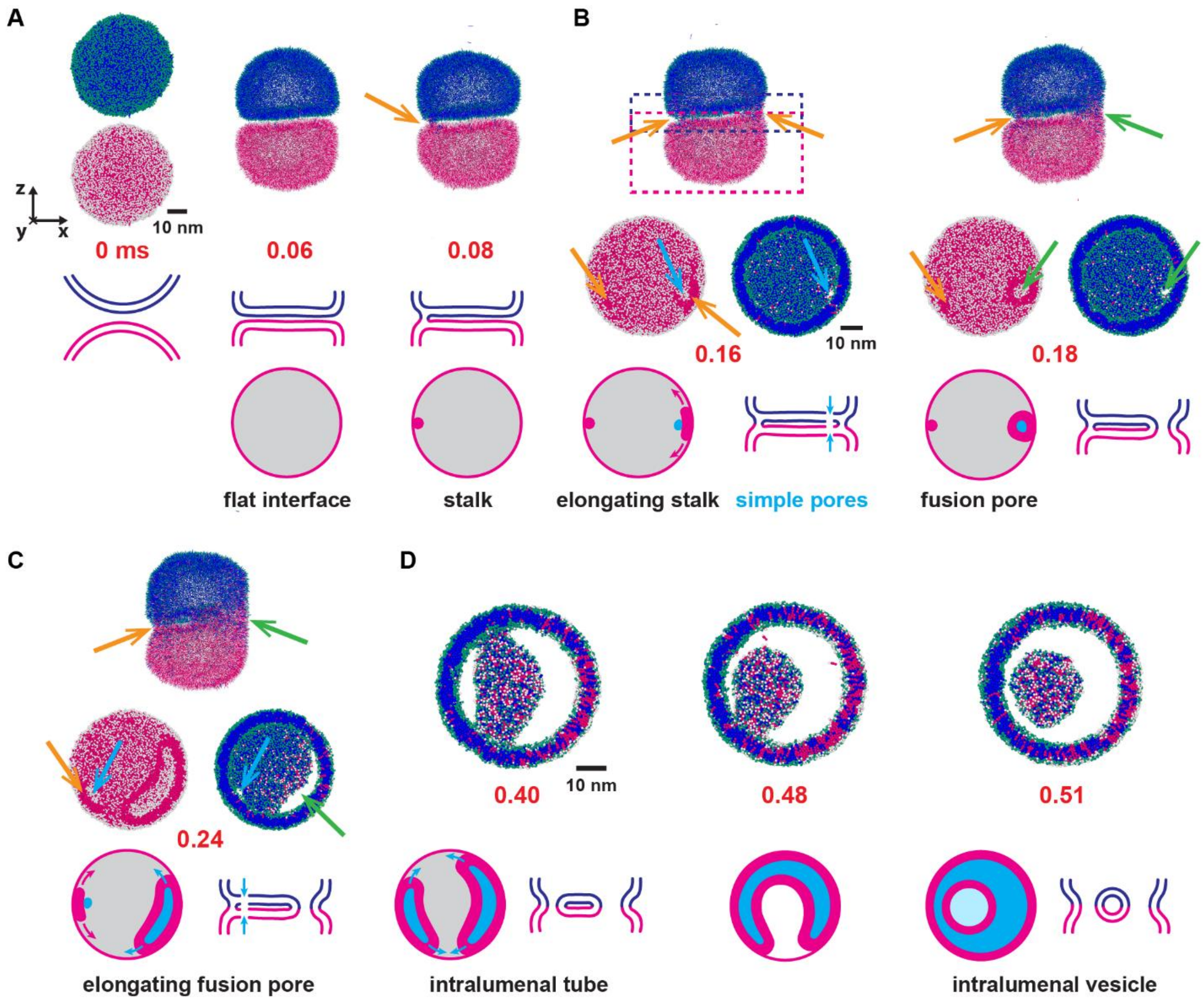


**Figure S2.** Detailed fusion pathway driven by brute force (same trajectory as in Fig. 3). Snapshots of two 50 nm vesicles pressed together by a 675 pN net force. Top vesicle lipids: white heads, magenta tails. Bottom vesicle lipids: green heads, blue tails. (A) A large flat membrane-membrane interface forms. Within 80 μs a hemifusion stalk (arrow) nucleates at the interface edge. (B) By 0.16 ms, a second stalk has nucleated and expanded along the interface edge (orange arrow, right) and a simple pore has formed in each vesicle (blue arrows) near the stalk's inner edge, transiently allowing contents leakage. By 0.18 ms the second stalk has encircled the simple pores, sealing them into a non-leaky fusion pore (green arrows). Middle row: orthographic projections of contents of the magenta or blue dashed boxes (magenta box includes only lipids belonging initially to lower vesicle). (C) The first stalk elongates and provokes a pair of simple pores (blue arrows). The fusion pore extends along the interface edge (green arrow). Middle row: orthographic projections as in (B). (D) Orthographic projections of blue dashed box contents as in (B). Two fusion pores expand along the interface edge, creating an intralumenal membrane tube. By 0.51 ms the fusion pores have merged, releasing an intralumenal vesicle.

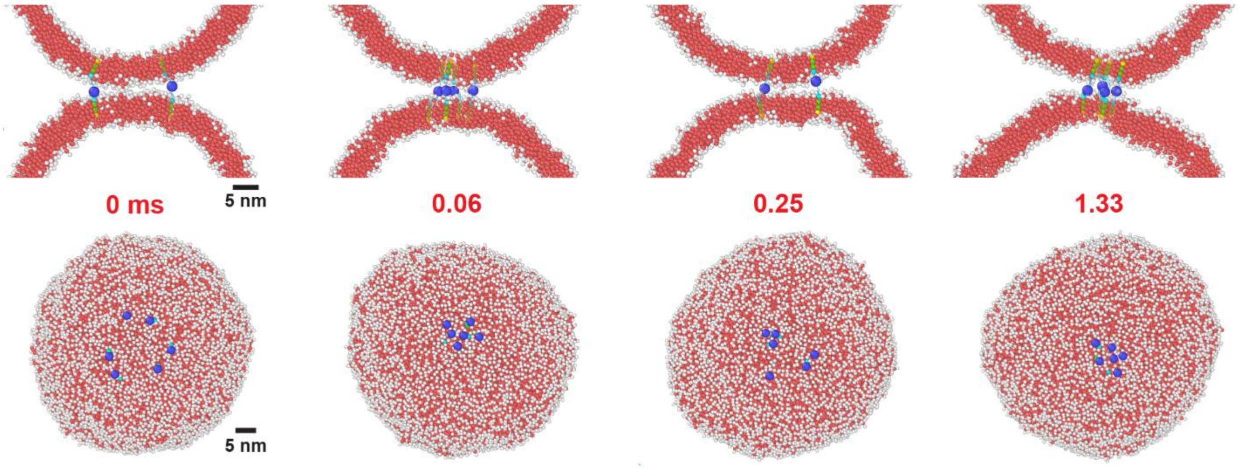


**Figure S3.** Non-rod-like complexes with a 2 nm diameter globular domain failed to hemifuse or fuse membranes in 20 runs each of ~ 1.4 ms. The six fusogens clustered between the vesicles within ~ 0.06 ms where they remained, except for occasional unclustered transients. Top row: ~ 5 nm thick cross-sections. Bottom row: top views of the lower vesicle.

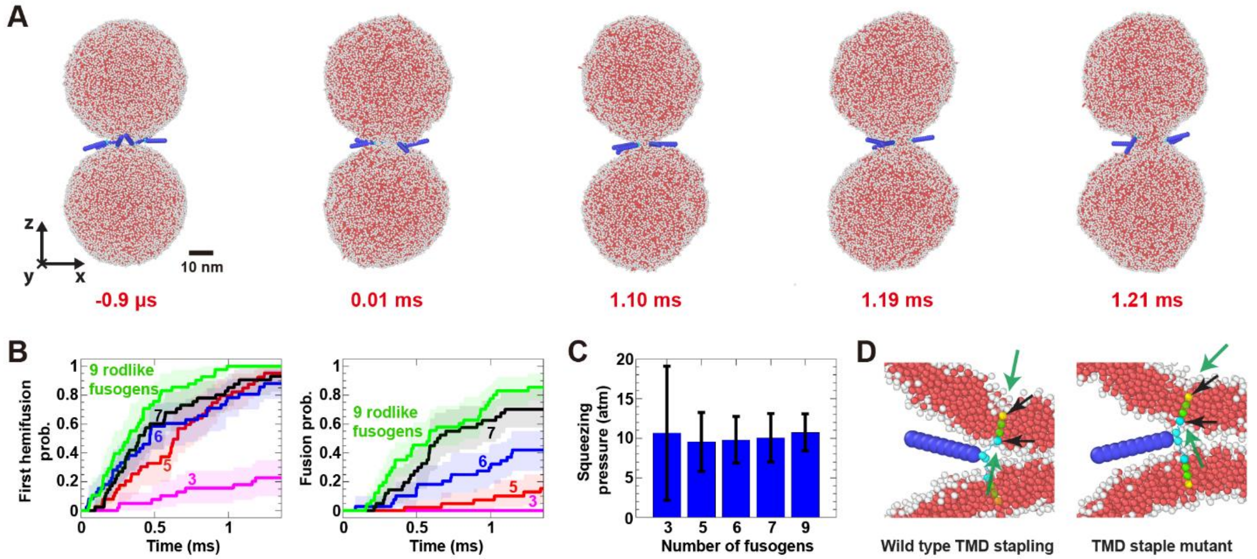


**Figure S4.** More rod-like fusogens promote faster fusion, and TMD-induced membrane thinning further catalyzes the process. (A) Side view of the fusion pathway driven by six model rod-like fusogens (same trajectory as in Fig. 4B). (B) Cumulative probability distributions of hemifusion and fusion times for different numbers of fusogens (n = 40 simulations for each). Shaded regions denote 95% confidence intervals computed via bootstrapping. (C) Total pressure squeezing the vesicle membranes together prior to hemifusion for different numbers of fusogens (n = 40 simulations for each). Error bars: SD. (D) Left: wild type (WT) rod-like fusogens with charged TMD staple beads (black arrows) induce local membrane thinning (green arrows). Right: rod-like fusogen mutants with dysfunctional neutral TMD staple beads (black arrows) fail to thin membranes (green arrows). No hemifusion or fusion occurred in 20 simulations with six mutant fusogens (~ 28 ms total simulation time).

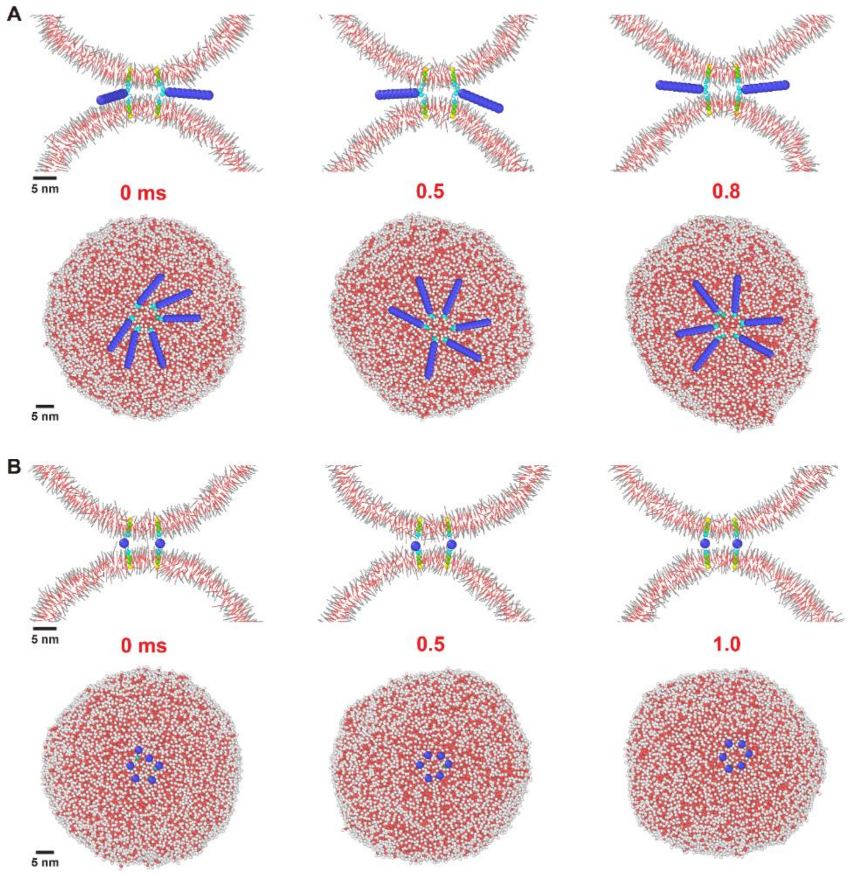


**Figure S5.** Snapshots from simulations with six rod-like (A) or non-rod-like (B) fusogens where the TMD positions were fixed in a ring of ~ 3 nm radius near the fusion site throughout the entire simulation.

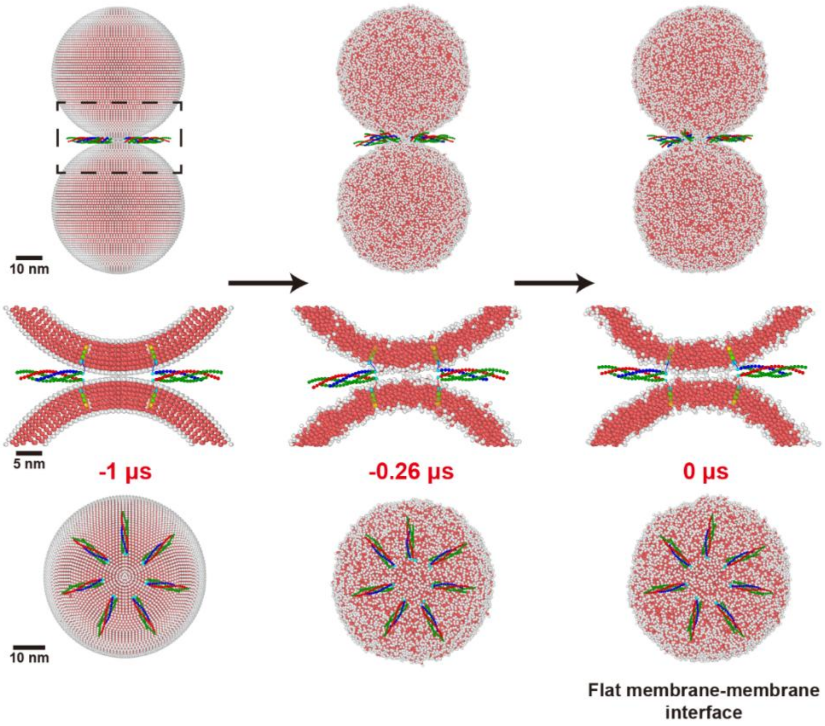


**Figure S6.** Snapshots from a simulation with seven SNAREs during a 1 μs equilibration period. The initial condition consisted of two vesicles with crystalized bilayers and no adhesion, bridged by seven SNARE complexes oriented radially in a ring whose center lies directly in the middle of the two vesicle centers. Entropic forces generated by the SNAREs maintained the fusion site clear, and within 1 μs, created a flat membrane-membrane interface with a radius of ~ 6 nm. Middle row shows cross-sectional blow-ups of the boxed region indicated top left.

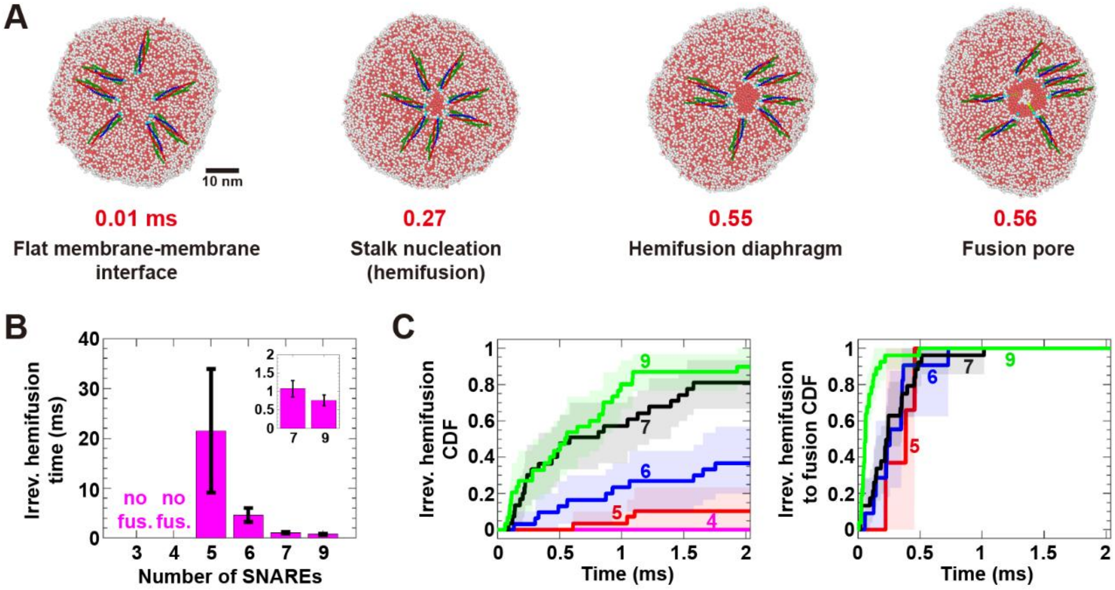


**Figure S7.** SNARE-mediated fusion pathway. (A) Additional perspective of the fusion pathway driven by seven SNAREs (same trajectory as in Fig. 5A). (B) Mean irreversible hemifusion time versus number of SNAREs (n = 30 simulations for each). Error bars: SEM. (C) Left: cumulative distributions of irreversible hemifusion times for different numbers of SNAREs (n = 30 smiulations for each). Right: cumulative distributions of irreversible hemifusion to fusion times from simulations where fusion occurred. Shaded regions denote 95% confidence intervals computed via bootstrapping.

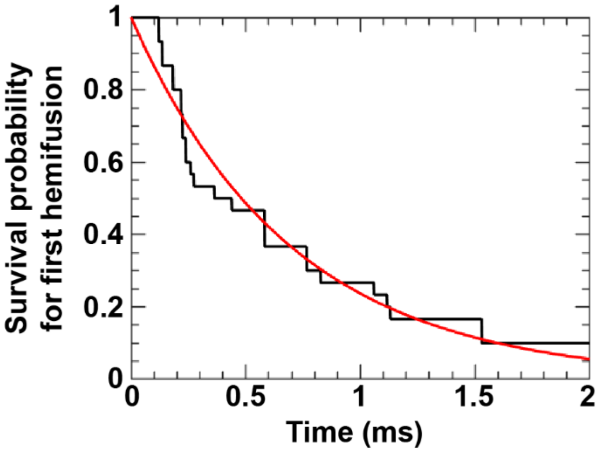


**Figure S8.** Waiting times for the first hemifusion event follow an exponential distribution. First hemifusion times from $n$ simulations were resampled with replacement to generate a dataset of $n$ times. A survival probability curve was constructed from these times (black) and fit to $e^{-t/\tau}$ (red), where $\tau$ is the estimated mean first hemifusion time. The example shown here corresponds to simulations with seven SNAREs (n = 30), yielding $\tau = 0.69\ \mathrm{ms}$. This resampling and fitting procedure was repeated 200 times to estimate the overall mean first hemifusion time. The same approach was used to estimate mean waiting times for all event types in simulations with SNAREs, EFF-1, and model rod-like fusogens.

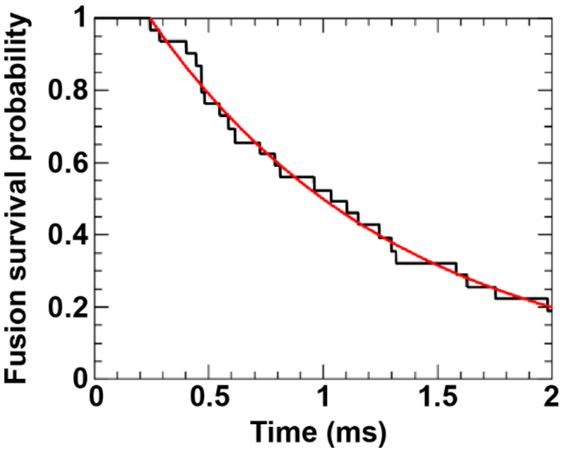


**Figure S9.** Fusion times follow an exponential distribution. The mean fusion survival probability curve (black) was obtained from 30 simulations with seven SNAREs by averaging over 200 bootstrap resamplings. The portion of the curve with $t \geq t_0$ was fit to $e^{-(t-t_0)/\tau}$ (red), where $t_0$ is the minimum fusion time across all simulations. The red fit was differentiated to obtained the fusion time probability distribution shown in Fig. 5I.

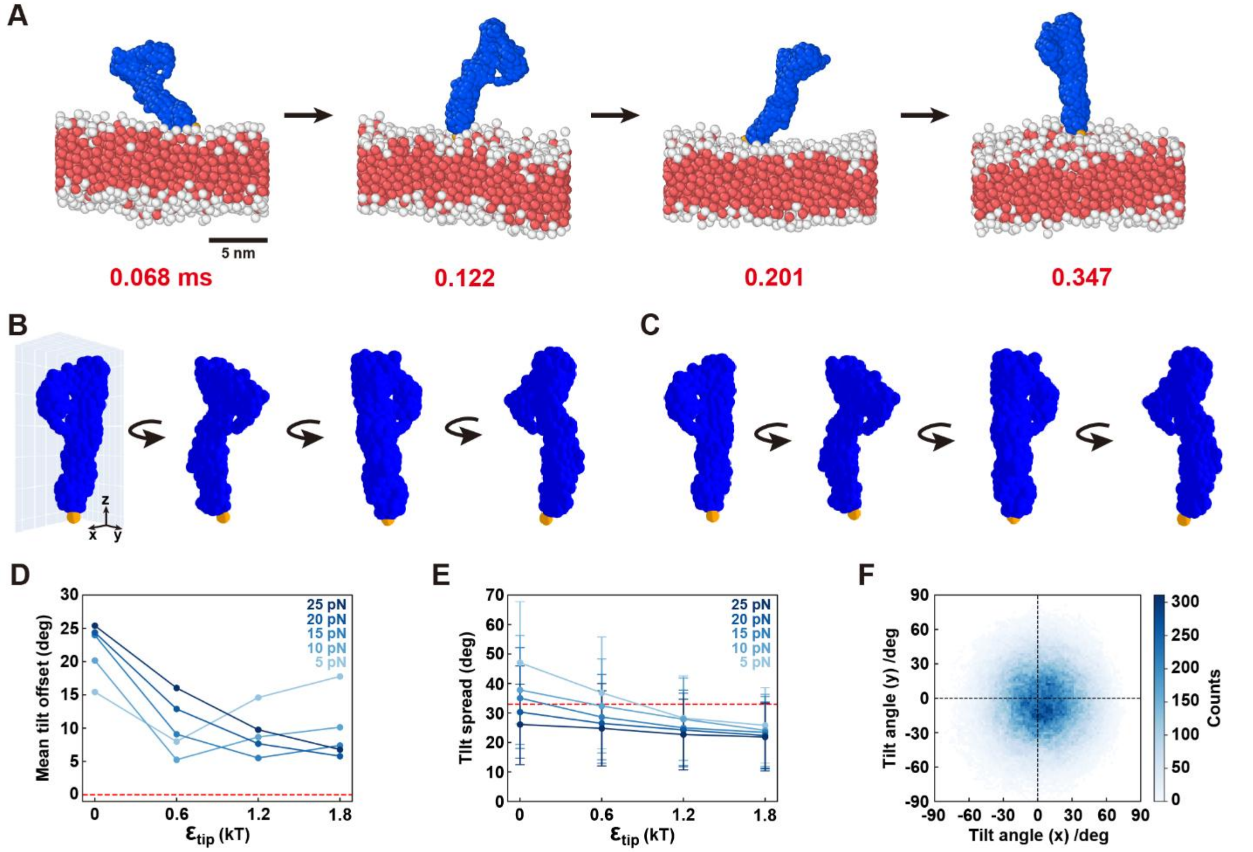


**Figure S10.** The orientation of EFF-1 monomers on the membrane is determined by the strength of the acidic tip. A single EFF-1 monomer was simulated with its TMD anchored in a planar membrane, and tethered to the TMD via a fixed-force spring. The tip-lipid head interaction energy well depth $\epsilon_{\text{tip}}$ was varied between 0-1.8 $k_BT$ and the fixed-force spring tension $f_{\text{LD}}$ was varied between 5-25 pN. (A) Snapshots from a simulation with $\epsilon_{\text{tip}} = 0.6\,k_BT,\ f_{\text{LD}} = 10\text{ pN}$. (B) Upright orientation of the EFF-1 monomer on the membrane, consistent with the upright orientation observed experimentally (3). Four views are shown, corresponding to 90° rotations about the membrane normal (z axis). The acidic tip is shown in orange. (C) Mean monomer orientation from simulations with $\epsilon_{\text{tip}} = 0.6\,k_BT$ and $f_{\text{LD}} = 10$ pN. (D) Mean tilt offset (i.e. tilt of the average monomer orientation relative to the upright reference shown in (B)) as a function of $\epsilon_{\text{tip}}$ and $f_{\text{LD}}$. (E) Tilt spread (i.e. mean deviation of the monomer from its average orientation) as a function of $\epsilon_{\text{tip}}$ and $f_{\text{LD}}$ (n ≈ 295,000 measurements per condition). Dashed red lines in (D) and (E) indicate the experimental mesurements reported in (3), where the mean orientation was upright (zero mean tilt offset) and the tilt spread was 33±17° relative to the membrane normal. Simulations with $\epsilon_{\text{tip}} = 0.6\,k_BT$ and $f_{\text{LD}} = 10$ pN best reproduced these experimental results. (F) Monomer tilt distribution from simulations with $\epsilon_{\text{tip}} = 0.6\,k_BT$ and $f_{\text{LD}} = 10$ pN. The distance from the origin to the distribution center indicates the mean tilt offset shown in (D), ~ 5°, while the mean distance of all observed configurations from the distribution center reflects the tilt spread shown in (E), 32±16°.

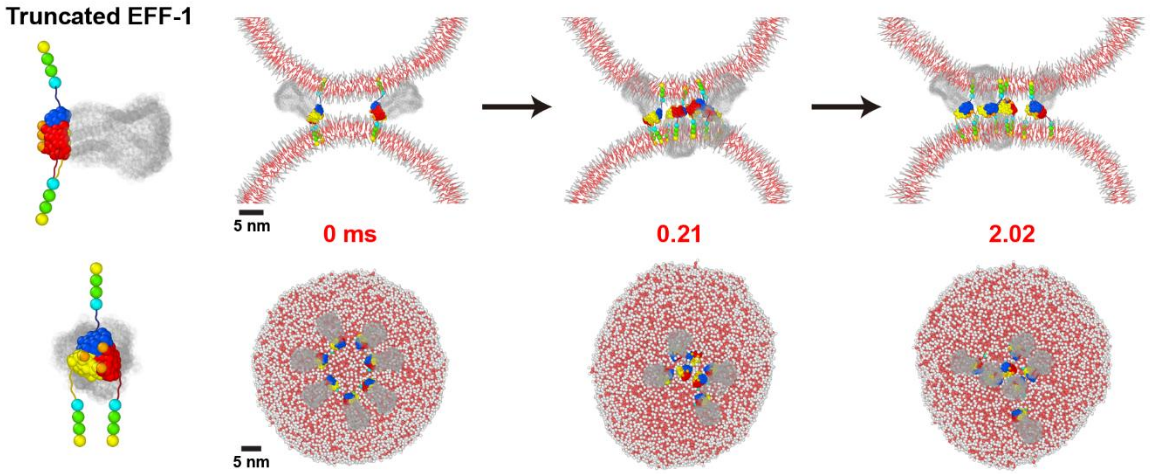


**Figure S11.** Non-rod-like EFF-1 mutants truncated to ~ 2.3 nm length (removed beads shown grey) failed to hemifuse or fuse membranes. Right: the mutants cluster between the vesicles and block hemifusion (seven fusogens, $\epsilon_{\text{tip}} = 1.2\ k_BT$).

**Table S1.** Model parameters

| Symbol | Meaning | Value | Legend |
|---|---|---|---|
| $\sigma$ | Length unit in the Cooke-Deserno model | 0.88 nm | |
| $\epsilon$ | Energy unit in the Cooke-Deserno model | 0.6 $k_B T$ | (A) |
| $m$ | Mass unit in the Cooke-Deserno model | $3.05 \times 10^{-25}$ kg | (A) |
| $D_{\mathrm{tail}}$ | Lipid tail bead diameter | $\sigma$ | (A) |
| $D_{\mathrm{head}}$ | Lipid head bead diameter | $0.95\sigma$ | (A) |
| $D_{\mathrm{rod}}$ | Rod-like fusogen bead diameter | 2 nm | (B) |
| $D_{\mathrm{TMD}}$ | TMD bead diameter | 1 nm | (B) |
| $L_{\mathrm{rod}}$ | Rod-like fusogen length | 10 nm | (B) |
| $D_{\mathrm{SNARE}}$ | SNARE bead diameter | 0.66 nm | |
| $D_{\mathrm{EFF-1}}$ | EFF-1 bead diameter | 0.88 nm | |
| $m_{\mathrm{tail}}$ | Mass of a lipid tail bead | $m$ | |
| $m_{\mathrm{head}}$ | Mass of a lipid head bead | $m$ | |
| $m_{\mathrm{rigid}}$ | Mass of each rigid body (fusogens and TMDs) | $m$ | (C) |
| $\gamma_{\mathrm{b}}$ | Drag coefficient of a lipid or gas bead | $\sqrt{m\epsilon}/\sigma$ | |
| $\gamma_{\mathrm{TMD}}$ | Drag coefficient of the TMD | $\gamma_{\mathrm{b}}$ | |
| $\gamma_{\mathrm{rod}}$ | Drag coefficient of the rod-like fusogen | $\gamma_{\mathrm{b}}$ | |
| $\gamma_{\mathrm{glob}}$ | Drag coefficient of the non-rod-like fusogen | $\gamma_{\mathrm{b}}$ | |
| $\gamma_{\mathrm{SNARE}}$ | Drag coefficient of the SNARE complex | 69 $\gamma_{\mathrm{b}}$ | (D) |
| $\gamma_{\mathrm{EFF-1}}$ | Drag coefficient of the EFF-1 fusogen | 630 $\gamma_{\mathrm{b}}$ | (D) |
| $\gamma_{\Delta\mathrm{EFF-1}}$ | Drag coefficient of the truncated EFF-1 fusogen | 76 $\gamma_{\mathrm{b}}$ | (D) |
| $\Delta t$ | Time step in all simulations except those with EFF-1 | 0.068 ns, 0.68 ps | (E) |
| $\Delta t_{\mathrm{EFF-1}}$ | Time step in EFF-1 simulations | 0.034 ns, 0.34 ps | (F) |

(A) Obtained from ref. (4).
(B) Approximated based on the dimensions of the SNARE complex, obtained from ref. (8).
(C) SNAREs, model rod-like fusogens, and TMDs have the moment of inertia matrix of a uniform density cylinder. Globular fusogens have the moment of inertia matrix of a uniform density sphere. The moment of inertia of EFF-1 was calculated using the atomic coordinates from the crystal structure (1).
(D) Calculated using the free-draining approximation, $\gamma_{\mathrm{fusogen}} = N\gamma_{\mathrm{b}}$, where $N$ is the number of beads used to represent the fusogen.
(E) A time step of 0.068 ns was used in simulations. During equilibration, the time step was 0.68 ps.
(F) A time step of 0.034 ns was used in simulations. During equilibration, the time step was 0.34 ps.

**Movie S1.** Top view of a simulated system (0-0.41 ms) where six non-rod-like, globular fusogens initiated in a ring rapidly gather near the fusion site, blocking hemifusion and fusion. The video captures the small fraction of lipids (~0.1%) that partition into solution from the membranes. These were omitted from figures for clarity.

**Movie S2.** Side view of a simulated system (0-0.41 ms) where six non-rod-like, globular fusogens initiated in a ring rapidly gather near the fusion site and remain there, blocking hemifusion and fusion.

**Movie S3.** Top view of a simulated system (0.84-1.28 ms) showing hemifusion and fusion driven by six rod-like fusogens.

**Movie S4.** Side view of a simulated system (0.84-1.28 ms) showing hemifusion and fusion driven by six rod-like fusogens.

**Movie S5.** Top view of a simulated system (0.14-1.07 ms) showing hemifusion and fusion driven by seven SNAREs.

**Movie S6.** Top view of a simulated system (0.14-0.54 ms) showing hemifusion and fusion driven by seven EFF-1 fusogens.

**Movie S7.** Top view of a simulated system (0-0.44 ms) where seven non-rod-like, truncated EFF-1 fusogens initiated in a ring gather unproductively near the fusion site and fail to drive hemifusion or fusion.
